# Task-Dependent Warping of Semantic Representations During Search for Visual Action Categories

**DOI:** 10.1101/2021.06.17.448789

**Authors:** Mo Shahdloo, Emin Çelik, Burcu A. Ürgen, Jack L. Gallant, Tolga Çukur

## Abstract

Object and action perception in cluttered dynamic natural scenes relies on efficient allocation of limited brain resources to prioritize the attended targets over distractors. It has been suggested that during visual search for objects, distributed semantic representation of hundreds of object categories is warped to expand the representation of targets. Yet, little is known about whether and where in the brain visual search for action categories modulates semantic representations. To address this fundamental question, we studied human brain activity recorded via functional magnetic resonance imaging while subjects viewed natural movies and searched for either *communication* or *locomotion* actions. We find that attention directed to action categories elicits tuning shifts that warp semantic representations broadly across neocortex, and that these shifts interact with intrinsic selectivity of cortical voxels for target actions. These results suggest that attention serves to facilitate task performance during social interactions by dynamically shifting semantic selectivity towards target actions, and that tuning shifts are a general feature of conceptual representations in the brain.

## Introduction

The ability to swiftly perceive the actions and intentions of others is a crucial skill for all social animals. In the human brain this ability has been attributed to a network of occipitotemporal, parietal and premotor areas collectively called the action observation network (AON) (Caspers et al., 2010; Molinari et al., 2013; Oberman et al., 2007; Rozzi and Fogassi, 2017). Recent reports suggest that the AON hierarchically represents diverse information pertaining to actions, ranging from shape and kinematics to action-effector interactions and action categories (Grafton and de C Hamilton, 2007; Handjaras et al., 2015; Oosterhof et al., 2010, 2012, 2013; Urgen et al., 2019; Wurm et al., 2017; Cavina-Pratesi et al., 2018). Low-level shape and movement kinematics are represented in occipitotemporal cortex and in the posterior bank of inferior temporal cortex (Jastorff and Orban, 2009). Effector type (e.g., foot, hand) is represented in ventral premotor cortex (Corbo and Orban, 2017; Jastorff et al., 2010), while parietal cortex represents higher-level action categories (Abdollahi et al., 2012; Ferri et al., 2015).

Several lines of evidence suggest that selective attention alters population responses to actions across this representational hierarchy. A host of electrophysiology (Muthukumaraswamy and Singh, 2008; Muthukumaraswamy et al., 2004; Puglisi et al., 2017, 2018; Schuch et al., 2010) and neuroimaging studies (Herrington et al., 2012; de Lange et al., 2008; Nicholson et al., 2017; Rowe et al., 2002; Safford et al., 2010) have examined effects of attention to relatively low-level action features. Using electroencephalography (EEG), Schuch et al. (2010) reported increased responses across the AON when attention was directed to action kinematics compared to a task-irrelevant visual feature (e.g., colour). Safford et al. (2010) presented overlapping moving tools and moving humans via simplified point-light displays (Johansson, 1973) and reported that attending to animate actors (i.e., humans) versus inanimate objects enhanced blood oxygen level dependent (BOLD) responses in superior temporal sulcus (STS). Nicholson et al. (2017) presented short action movies in controlled scenes and subjects attended to the manipulated objects or to action goals in different runs. Responses were enhanced in inferior frontal gyrus (IFG), occipitotemporal cortex, and middle frontal gyrus (MFG) as a result of attention to action goals, whereas attending to manipulated objects increased responses in parietal cortex and fusiform gyrus. In contrast, only few previous reports have further investigated the effects of attention to higher-level action features (Nastase et al., 2017, 2018). In a recent study, Nastase et al. (2017) presented movie clips of animals from five taxonomies (primates, ungulates, birds, reptiles, and insects) performing actions from four categories (eating, fighting, running, and swimming), and asked participants to attend either to taxonomy or action of the stimulus. Using searchlight analysis on representational dissimilarity matrices (RDMs) (Haxby et al., 2014, 2020a; Kriegeskorte and Kievit, 2013; Kriegeskorte et al., 2008; Nili et al., 2014), they reported that attending to performed actions alters multi-variate response patterns across anterior intraparietal sulcus (IPS) and premotor cortex.

Taken together, current electrophysiology and neuroimaging findings on selective attention to visual actions suggest that attention increases AON responses to target features ranging from action kinematics and goals to actors. That said, high-level semantic representations during targeted visual search for specific action categories remain understudied. Furthermore, prior studies did not question whether attending to action features causes simple baseline and gain changes in population responses, or rather elicits dynamic tuning shifts that can alter cortical representation. Recent evidence indicates that visual search for object categories shifts single-voxel category tuning toward target objects (Çukur et al., 2013). Therefore, it is likely that attention to action categories also causes tuning shifts to facilitate visual search.

Here we hypothesised that natural visual search for action categories induces semantic tuning shifts in single cortical voxels to expand the representation of target actions while compressing the representation of behaviourally irrelevant actions (Fig. 1). To test the tuning-shift hypothesis, we recorded whole-brain BOLD responses while human subjects viewed 60min of natural movies and covertly searched for either 15 *communication* actions or 26 *locomotion* actions among 109 action categories in the movies (see *Supplementary Methods*). Using spatially informed voxelwise modelling (Çelik et al., 2019), we measured category responses for hundreds of objects and actions in the movies separately for each individual subject and for each search task. We estimated a semantic space underlying action-category responses, and semantic tuning for action categories were measured by projecting voxel-wise model weights onto this space. Finally, semantic tuning profiles during the two search tasks were compared to quantify the magnitude and direction of tuning shifts in single voxels.

**Figure 1.**
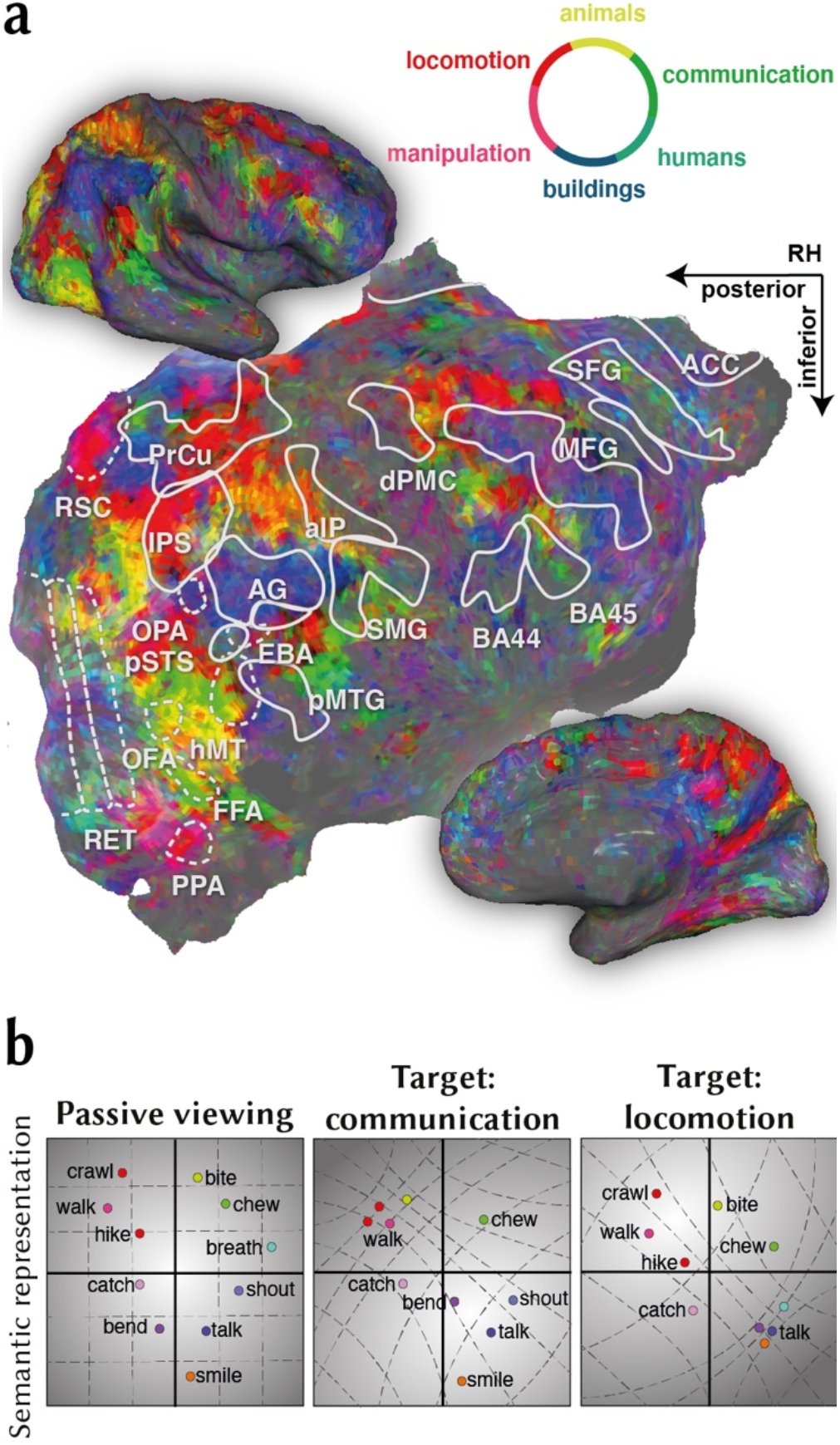
Hypothesised changes in semantic representation of action categories. Recent evidence suggests that the human brain organises hundreds of object and action categories in a semantic space that is distributed systematically across the cerebral cortex (Huth et al., 2012). **a.** Semantic representation for a single subject from Çukur et al. (2013) is shown on flattened cortical surface and on inflated hemispheres. Colours indicate tuning for different object or action categories (see colour legend). Regions of interest identified using conventional functional localizers are denoted by white borders. Abbreviations for regions of interest are listed in *Supplementary Methods*. **b.** In the semantic space, action categories that are semantically similar to each other are mapped to nearby points and semantically dissimilar actions are mapped to distant points. There is evidence that visual search for object categories warps semantic representation in favour of the targets by shifting single-voxel tuning for object categories toward target objects (Çukur et al., 2013). Thus, we hypothesised that visual search for a given action category should similarly expand the semantic representation of the target and semantically similar categories.

## Results

### Visual search modulates category responses

Little is known on whether and where in the brain natural visual search for action categories warps semantic representations. To answer this question, we investigated voxel-wise tuning for hundreds of object and action categories across cortex. Human subjects viewed natural movies and covertly searched for *communication* or *locomotion* actions. Category regressors were constructed to label presence of 922 distinct object and action categories in the movies. Separate category models were then fit in each voxel for each search task. These models enabled us to measure single-voxel category responses during each search task (Fig. 2a, see *Experimental Procedures*).

**Figure 2.**
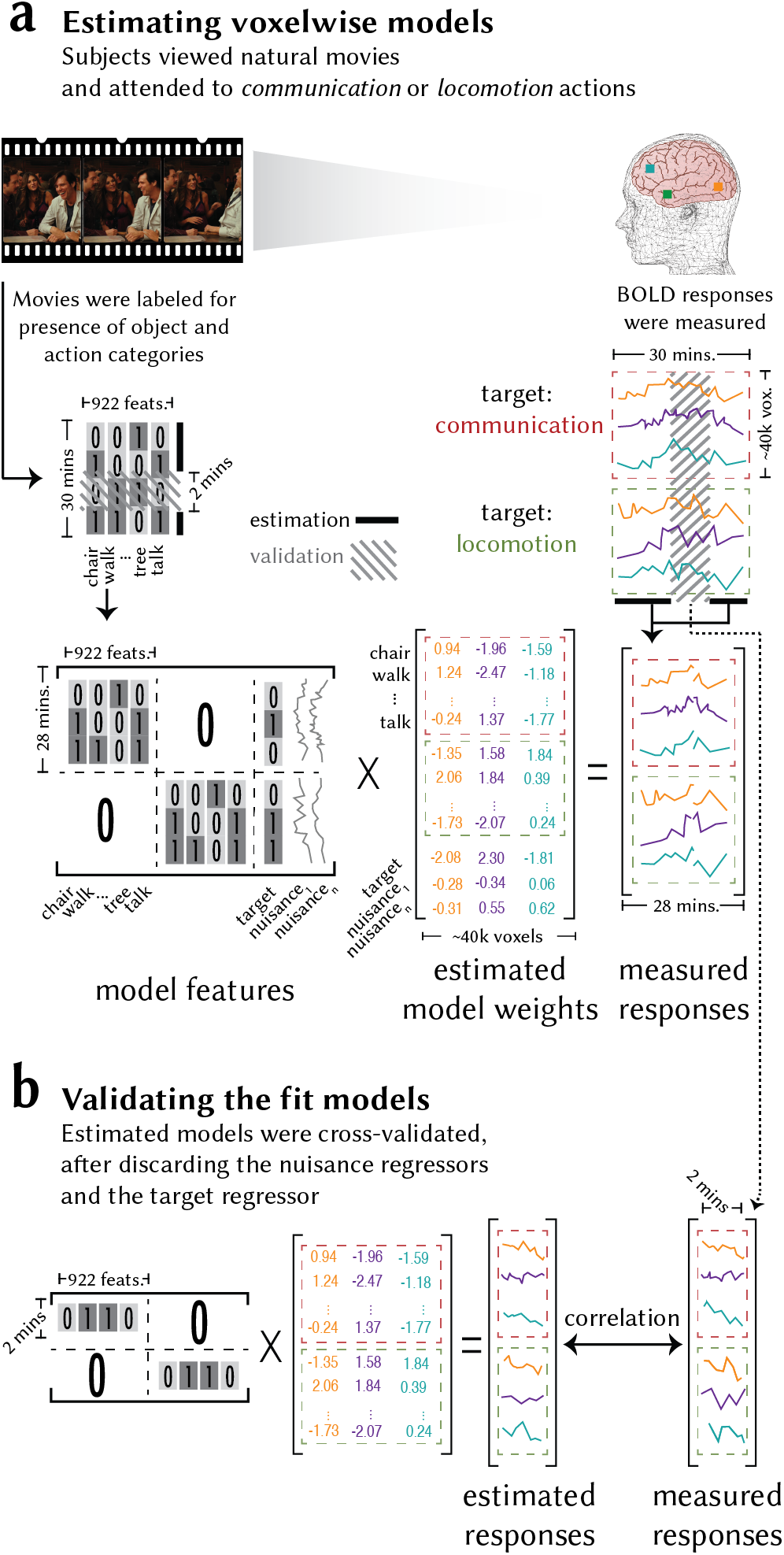
Model fitting and validation procedure. Undergoing fMRI, human subjects viewed 60mins of natural movies and covertly searched for *communication* or *locomotion* action categories while fixating on a central dot. **a.** An indicator matrix was constructed that identified the presence of each of the 922 object and action categories in each 1-sec clip of the movies. Nuisance regressors were included to account for head-motion, physiological noise, and eye-movement confounds. An additional nuisance regressor was included to account for target detection confounds. In a nested cross-validation (CV) procedure, regularized linear regression was used to estimate separate category model weights (i.e., category responses) for each search task that mapped each category feature to the recorded BOLD responses in single voxels. **b.** Accuracy of the fit models was cross-validated by measuring prediction performance on the held-out data in each CV fold, after discarding the nuisance regressors and the target regressor. Prediction score of the fit models was taken as product-moment correlation coefficient between estimated and measured BOLD responses, averaged across the two search tasks.

As natural stimuli contain correlations among various levels of features, there is a possibility that estimated category responses are confounded by voxel tuning for low- and intermediate-level scene features. To rule out this potential confound, we measured the response variance explained by low-level motion-energy features, and intermediate-level spatiotemporal interest point (STIP) features. Motion-energy features were constructed using a pyramid of spatiotemporal Gabor filters (Nishimoto and Gallant, 2011). STIP features, providing an intermediate representational basis for human actions, were constructed by measuring optical flow over interest points with significant spatiotemporal variation (Laptev et al., 2008). We identified voxels in which the category model explained unique response variance after accounting for these alternative features via variance partitioning, and subsequent analyses were conducted on this set of uniquely explained voxels. To prevent bias in voxel selection due to attention, variance partitioning was performed on a separate dataset collected for this purpose (i.e., *passive-viewing dataset*, see *Experimental Procedures*). We find that the category model explains unique response variance after accounting for low- and intermediate-level features in 39.6±0.4% of cortical voxels (mean±sem across five subjects; bootstrap test, *q*(FDR)<0.05; Supp. Figs. 1, 2), yielding 12,500-17,091 voxels in individual subjects (henceforth called *the semantic voxels*).

Comparison of estimated category responses across search tasks would be justified only if the fit models can accurately predict BOLD responses that were held-out during model fitting. To assess prediction performance of the fit category models, we measured average prediction scores across the two search tasks, taken as product-moment correlation coefficient between the predicted and measured held-out responses (Fig. 2b). Category models have high prediction scores (greater than 1 std above the mean) in 38.8±0.1% of the semantic voxels. These include many voxels spread across the AON comprising occipitotemporal, parietal, and premotor cortices, as well as voxels in prefrontal and cingulate cortices (Fig. 3).

**Figure 3.**
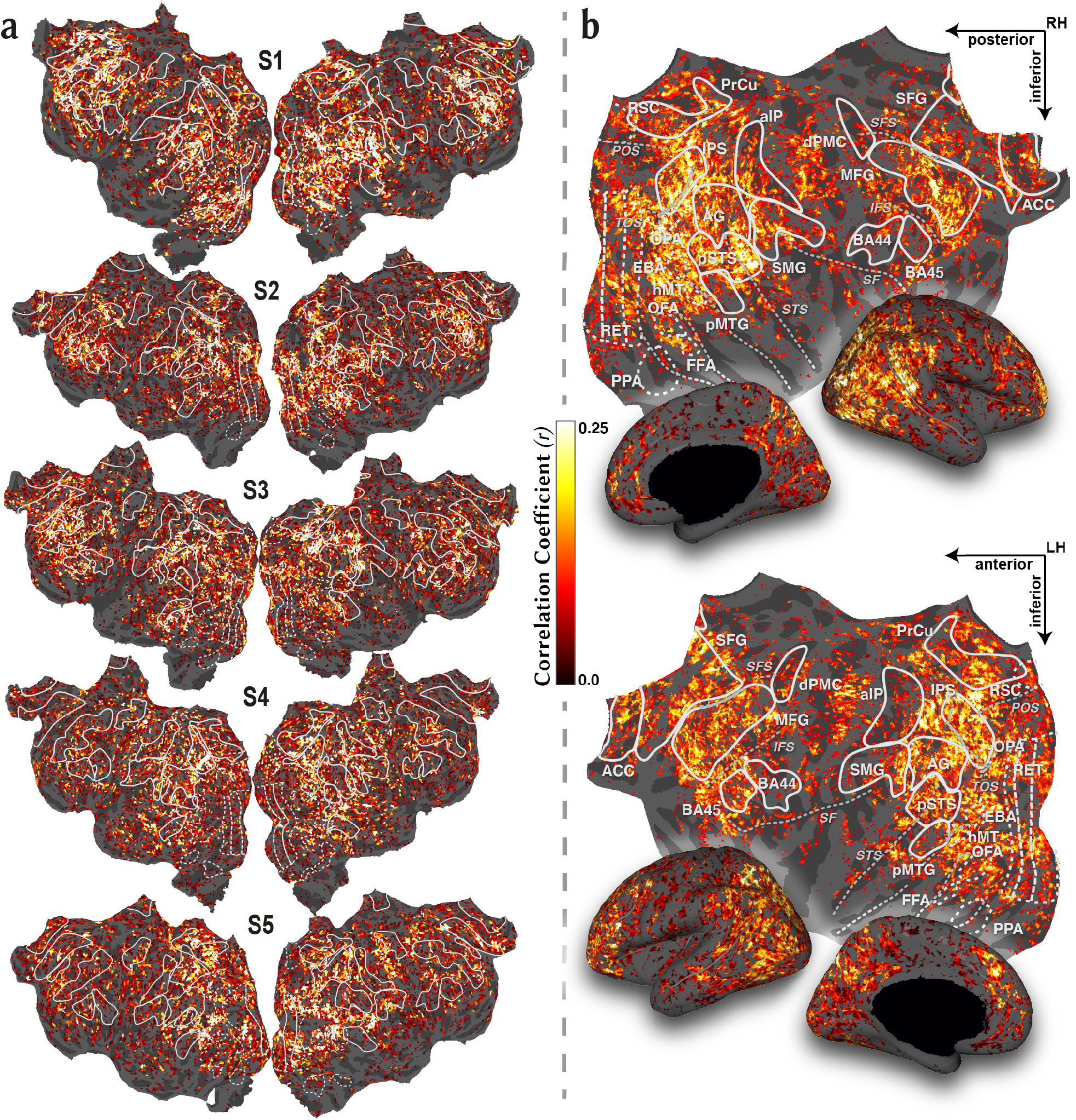
Prediction performance of the category model. To test the performance of fit category models, prediction score was calculated on held-out data as the product-moment correlation coefficient between the predicted category responses and measured BOLD responses, and it was averaged across the two search tasks. **a.** Prediction scores of the category model are plotted on flattened cortical surfaces of individual subjects. A variance partitioning analysis was used to quantify the response variance that was uniquely predicted by the category model after accounting for low- and intermediate-level stimulus features (see *Supplementary Methods*). Voxels where the category model did not explain unique response variance after accounting for these features were masked (bootstrap test, *q*(FDR)<0.05). **b.** To visualise single-subject results in a common space, prediction score values are shown following projection onto the standard brain template from Freesurfer and averaging across subjects. Regions of interest are illustrated by white borders. Several important sulci are illustrated by dashed grey lines. Abbreviations for regions of interest and sulci are listed in *Supplementary Methods*. The category model predicts responses across ventral-temporal, parietal, and frontal cortices well, suggesting that visual categories are broadly represented across visual and nonvisual cortex. Results can be explored via an interactive brain viewer at http://www.icon.bilkent.edu.tr/brainviewer/shahdloo_etal/.

A recent study provided the first evidence that attention can alter single-voxel category tuning profiles during search for object categories (Çukur et al., 2013). We thus hypothesised that visual search for action categories can also cause changes in voxel-wise category tuning. If attentional tuning changes are significant, the category models fit to individual search tasks should yield higher prediction scores than a null model fit by pooling data across the two search tasks. To test this prediction, we compared the prediction scores obtained from the category and null models. We find that the category model significantly outperforms the null model in 44.2±1.6% of semantic voxels (bootstrap test, *q*(FDR)<0.05). Additional control analyses further ensured that these attentional changes cannot be attributed to residual eye-movements, head-motion, physiological noise, or target-detection biases (see *Supplementary Methods*). Taken together, these results suggest that many cortical voxels in occipitotemporal, parietal, and prefrontal cortices encode high-level category information, and that action-based visual search significantly modulates category responses in single voxels.

### Visual search warps semantic representation of actions

Previous studies suggest that the human brain represents visual categories by embedding them in a continuous semantic space (Huth et al., 2012). To derive a semantic space underlying action category representations, we performed principal component analysis (PCA) on the model weights for action categories. Visual search for actions alters category model weights as reported here, so performing PCA on data from search tasks can bias estimates of the semantic space. Instead, we derived the semantic space using the passive-viewing dataset. Action categories that are semantically close to each other should project to nearby points in this space, whereas semantically dissimilar categories should project to distant points. The top twelve principal components (PCs) that explained more than 95% of the variance in responses were selected, which showed a high degree of inter-subject consistency (r=0.52±0.02 mean±sem across subjects; Supp. Fig. 3). To visually examine the semantic information captured by this space, we projected action categories onto the PCs (Fig. 4a; projections onto the first three dimensions that accounted for 72.8% of the response variance is shown in Supp. Fig. 4; loadings for all PCs are shown in Supp. Fig. 5). The first dimension seems to distinguish between self-movements (e.g., chew, yawn, eat) and actions that are targeted toward other humans or objects (e.g., reach, touch, drive). The second dimension seems to distinguish between dynamic versus static actions (e.g., raise, propel, dance versus breath, view). The third dimension appears to distinguish between actions that involve humans (e.g., communicate, reach, crouch) and dynamic actions (e.g., drive, fly, drag). These observations suggest that the estimated semantic space captures reasonable semantic variance across action categories in natural movies.

**Figure 4.**
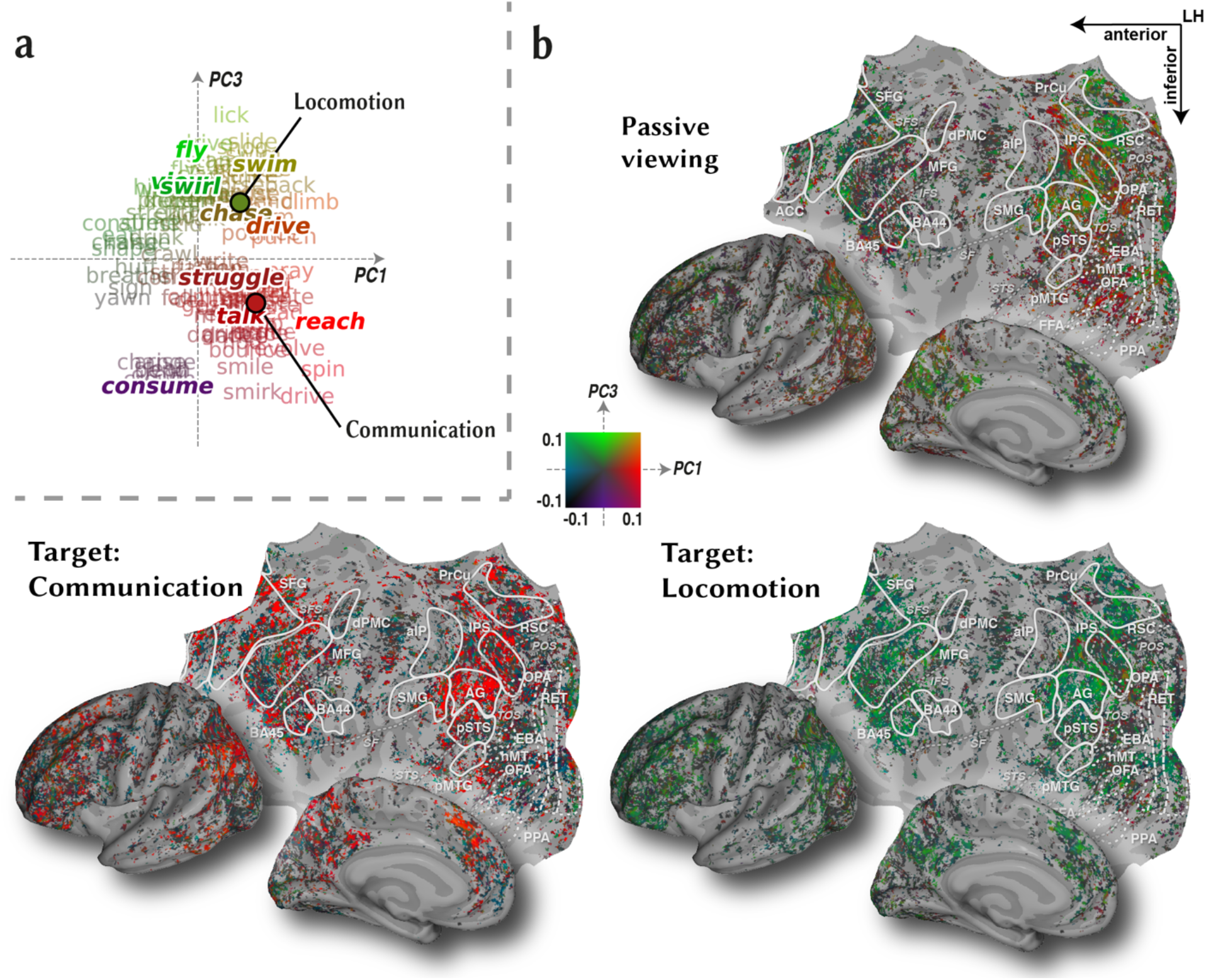
Attention warps semantic representation of action categories. To assess attentional changes, we projected voxel-wise tuning profiles onto a continuous semantic space. **a.** The semantic space was derived from principal components analysis (PCA) of tuning vectors measured during a separate passive-viewing task. To illustrate the semantic information embedded within this space, action categories were projected onto PC1 and PC3 that best delineate the target actions (Supp. Fig. 6; words in regular font show projections of individual categories). To facilitate illustration, categories were collapsed into 10 clusters and cluster centres were also projected onto the PCs (bold-italic words; see *Supplementary Methods*). Average location of the *communication* and *locomotion* actions are specified with red and green dots. **b.** Action category responses during passive viewing and during the two search tasks were projected onto the semantic space, and a two-dimensional colourmap was used to colour each voxel based on the projection values along PC1 and PC3 (see legend). Projections in individual subjects were mapped onto the standard brain template from Freesurfer, and average projections across subjects are displayed. Figure formatting is identical to Fig. 3. Many voxels across occipitotemporal, parietal, and prefrontal cortices shift their tuning toward targets, suggesting that attention warps semantic representations of actions. Specifically, voxels in inferior posterior parietal cortex, cingulate cortex, and anterior inferior prefrontal cortex shift their tuning toward *communication* during search for *communication* actions. Meanwhile, voxels in superior posterior and medial parietal cortex shift their tuning toward *locomotion* during search for *locomotion* actions. Results can be explored via an interactive brain viewer at http://www.icon.bilkent.edu.tr/brainviewer/shahdloo_etal/.

Previous evidence suggests that visual search shifts single-voxel tuning profiles to expand the representation of the targets (Çukur et al., 2013). Thus, it is possible that action-based visual search also shifts semantic tuning in single-voxels towards the target category. To investigate this possibility, we projected action category responses onto the semantic space. The first and third PCs maximally differentiated between actions belonging to the target categories (i.e., *communication* versus *locomotion* categories, Supp. Fig. 6). Therefore, we visually compared the projections onto these PCs across the two search tasks. We observe that attention causes semantic tuning modulations broadly across cortex (Fig. 4b; see Supp. Figs. 7-11 for results in individual brain spaces). Specifically, voxels in inferior posterior parietal cortex (PPC), cingulate cortex, and anterior inferior prefrontal cortex shift their tuning toward communication during search for communication actions. Meanwhile, voxels in superior PPC and medial parietal cortex shift their tuning toward locomotion during search for locomotion actions. Several reports suggest involvement of superior inferior PPC in representing locomotion actions (Corbo and Orban, 2017), and inferior PPC in representing communication actions (Abdollahi et al., 2012, Rizzolatti and Matelli, 2003). Therefore, our findings suggest that during search for a given action category, tuning shifts toward the target category are most prominent in voxels that are primarily selective for the target.

### Visual search for action categories shifts single-voxel semantic tuning profiles

Our inspection of semantic representations during visual search reveals that attention broadly modulates high-level action representations by shifting semantic tuning profiles in single voxels. To quantify the magnitude and direction of these tuning changes, we separately measured semantic selectivity for *communication* and *locomotion* action categories in each search task. For each voxel, a tuning shift index (TSI_all_∈ [-1,1]) was taken as the difference in semantic selectivity for targets when they were attended versus unattended. A positive TSI_all_ indicates shifts towards the target, a negative TSI_all_ indicates shifts away from the target, and a TSI_all_ of 0 suggests no change in between tasks (see *Experimental Procedures*). We find that voxels across many cortical regions shift their tuning toward the attended category (Fig. 5a; see Supp. Figs. 12a-16a for results in individual brain spaces). Figure 7a shows respective tuning shifts in relevant regions of interest (ROIs). Tuning shifts are significantly greater than zero in many areas across AON including occipitotemporal cortex (posterior STS, pSTS; posterior MTG, pMTG), posterior parietal cortex (intraparietal sulcus, IPS; AG, SMG), and premotor cortex (Brodmann’s areas 44, 45, BA44/45; bootstrap test *p*<0.05; Fig. 7a). This result suggests that focused attention to specific action categories shifts semantic tuning toward targets in single-voxels, and that these attentional modulations are present at all levels of the AON hierarchy including occipitotemporal cortex. In contrast, brain areas that are not specifically selective for actions (Kilintari et al., 2014; Nelissen et al., 2006, 2011) –such as the low-level motion-selective area (human middle temporal area, hMT) and the extrastriate body area (EBA) that is selective for human body-parts– do not exhibit significant shifts in selectivity for actions (*p*>0.05).

**Figure 5.**
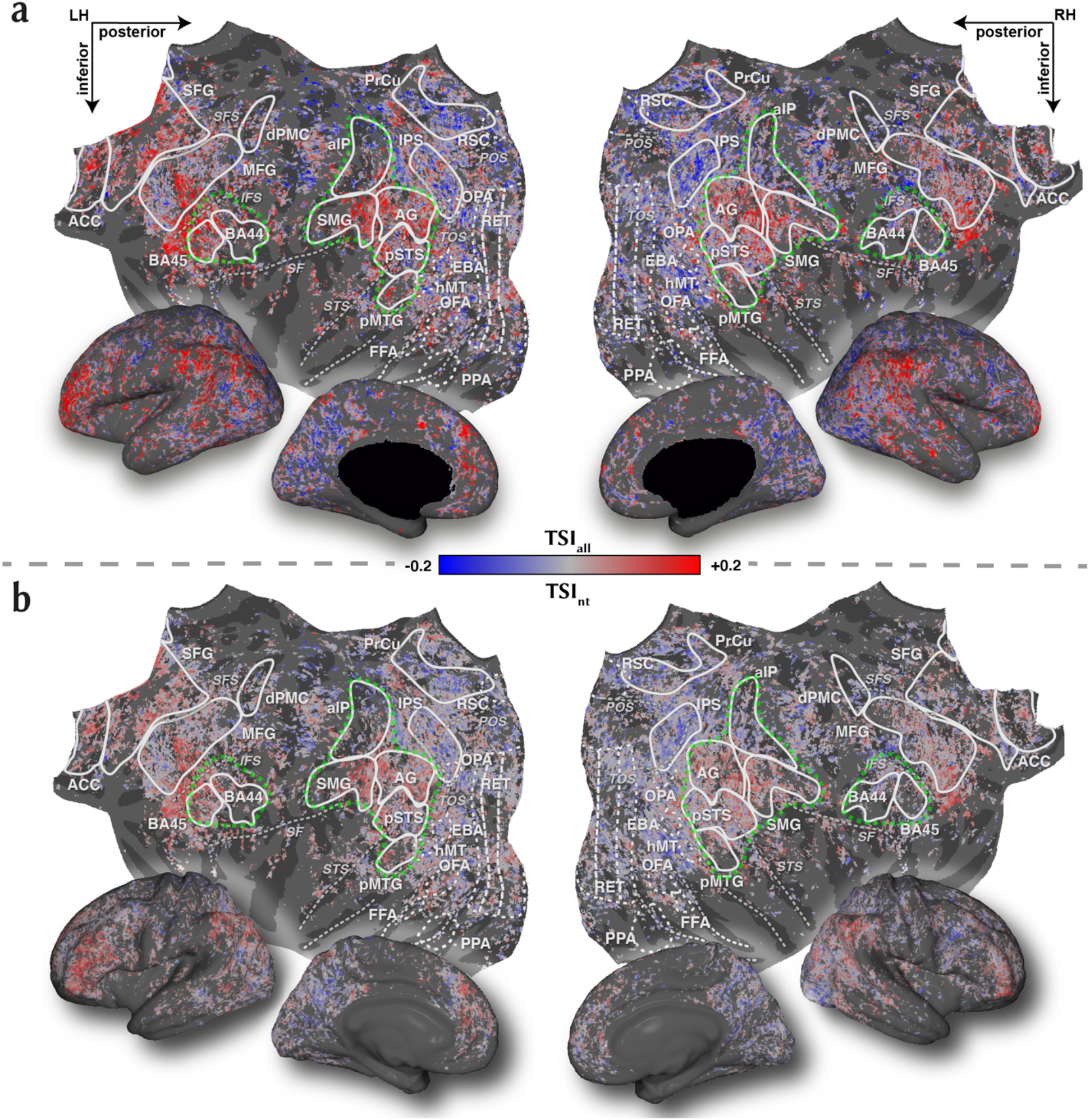
Cortical distribution of tuning shifts. **a.** To quantify the tuning shifts for the attended versus unattended categories, a tuning shift index (TSI_all_∈[-1,1]) was calculated for each voxel. Tuning shifts toward the attended category would yield positive TSI (red colour), whereas negative TSI would indicate shifts away from the attended category (blue colour). TSI_all_ values from individual subjects were projected onto the standard brain template and averaged across subjects. Figure formatting is identical to Fig. 3. AON is outlined by green dashed lines. Voxels across many cortical regions shifted their tuning toward the attended category. These include regions across AON (occipitotemporal cortex, posterior parietal cortex, and premotor cortex), lateral prefrontal cortex, and anterior cingulate cortex. **b.** To examine how representation of nontarget action categories changes during visual search, we measured a separate tuning shift index specifically for these categories (TSI_nt_). TSI_nt_ values from individual subjects were projected onto the standard brain template and averaged across subjects. TSI_nt_ shows a similar distribution to TSI_all_ shown in **a**, albeit with lower magnitude. Tuning shift for nontarget categories is positive across many voxels within posterior parietal cortex and anterior prefrontal cortex, suggesting a more flexible semantic representation of actions in these cortices, compared to occipitotemporal AON nodes. Results can be explored via an interactive brain viewer at http://www.icon.bilkent.edu.tr/brainviewer/shahdloo_etal/.

Prior evidence suggests that during category-based visual search, semantic tuning shifts grow stronger n toward later stages of semantic processing (Çukur et al., 2013). Here, we find that semantic tuning shifts in AG and SMG are significantly stronger than those in occipitotemporal (pSTS, pMTG) and premotor cortices (dorsal premotor cortex, dPMC; BA44/45; *p*<0.05). Therefore, the tuning shifts reported here could indicate that AG and SMG are higher nodes in the hierarchy of semantic representation of action categories. In a previous study, we reported that in medial prefrontal cortex visual search for object categories causes tuning shifts toward targets while it causes tuning shifts away from targets in voxels in precuneous (PrCu) and temporo-parietal junction (TPJ; Çukur et al., 2013). Similarly, here we find that visual search for action categories causes negative tuning shifts in many voxels across PrCu and TPJ. These results suggest that these areas might be involved in distractor detection and in error monitoring during visual search for actions (Corbetta and Shulman, 2002).

### Visual search shifts semantic tuning for nontarget action categories

Natural visual search for object categories was previously suggested to cause changes in representations of not only targets but also nontarget categories (Çukur et al., 2013; Seidl et al., 2012). Thus, it is likely that action-based visual search shifts semantic tuning for nontarget categories. To address this important question, we first examined the separate contributions of tuning changes for target versus nontarget categories to the overall tuning shifts. Specifically, we measured the fraction of overall tuning shifts that can be attributed to the target categories versus nontarget categories (i.e., all categories excluding *communication* and *locomotion* actions). We find that both target and nontarget categories significantly contribute to the overall tuning shifts (bootstrap test, *p*<0.05). However, as would be expected, target categories account for a relatively larger fraction of the overall tuning shifts compared to nontarget categories in all studied ROIs, except in early visual cortex (*p*<0.05; Supp. Fig. 17). Next, to explicitly quantify tuning shifts for nontarget categories, we calculated a separate tuning shift index exclusively on nontarget categories (TSI_nt_). To calculate TSI_nt_, the 109-dimensional action category response vectors were masked to select nontarget categories, prior to projection onto the semantic space (see *Experimental Procedures*). We observe that tuning shift for nontarget categories is generally smaller than the overall tuning shift (Fig. 5b versus Fig. 5a; see Supp. Figs. 12b-16b for results in individual brain spaces). Yet, TSI_nt_ is significant in AG, SMG, and BA45 (*p*<0.05; Fig. 7b). These results suggest that, compared to occipitotemporal areas, attention more diversely warps semantic representations in parietal and premotor AON nodes by shifting tuning for both target and nontarget categories.

### Tuning shifts interact with intrinsic selectivity of cortical voxels for action categories

A recent study on visual attention has reported that in strongly object-selective regions voxel tuning for a preferred object might be robust against attention directed to a nonpreferred object (e.g., *houses* for fusiform face area, FFA, and *faces* for parahippocampal place area, PPA; Çukur et al., 2013). This previous result suggests that the degree of response modulations in a brain region might depend on the alignment between the search target and the intrinsically preferred object. It is thus likely that tuning shifts during search for an action category also interact with the intrinsic selectivity of cortical voxels for the target category. Tuning shifts as measured by TSI signal an overall increase in relative selectivity for target versus nontarget categories, aggregated across search tasks. Yet, interaction of tuning shifts with intrinsic selectivity for action categories is task-specific by definition. Therefore, to examine potential interactions, we calculated a target preference index (PI∈[-1,1]) separately during search for communication actions (PI_com_) and during search for locomotion actions (PI_loc_). PI_com_ was taken as the difference in selectivity for *communication* versus *locomotion*, during search for communication actions. Analogously, PI_loc_ was taken as the difference in selectivity for *locomotion* versus *communication*, during search for locomotion actions.

Voxel-wise PI_com_ and PI_loc_ values were projected onto cortical flat maps for visual inspection (Fig. 6; see Supp. Figs. 12c-16c for results in individual brain spaces) and quantitatively examined in ROIs (Fig. 7c, d). We observe that semantic tuning in areas with indiscriminate selectivity for behaviourally relevant action categories (e.g., selective for low-level visual features or static object categories) show insignificant shifts regardless of the search task. Meanwhile, many voxels across anterior parietal, occipital, and cingulate cortices –with intrinsic action category preferences– show differential preference for one of the two target action categories as indicated by high PI index during either search for communication or search for locomotion actions. Lastly, semantic tuning in voxels across posterior parietal and anterior prefrontal cortices with broad selectivity for actions shift toward the attended category irrespective of the search target. These specific cases are discussed in detail below.

**Figure 6.**
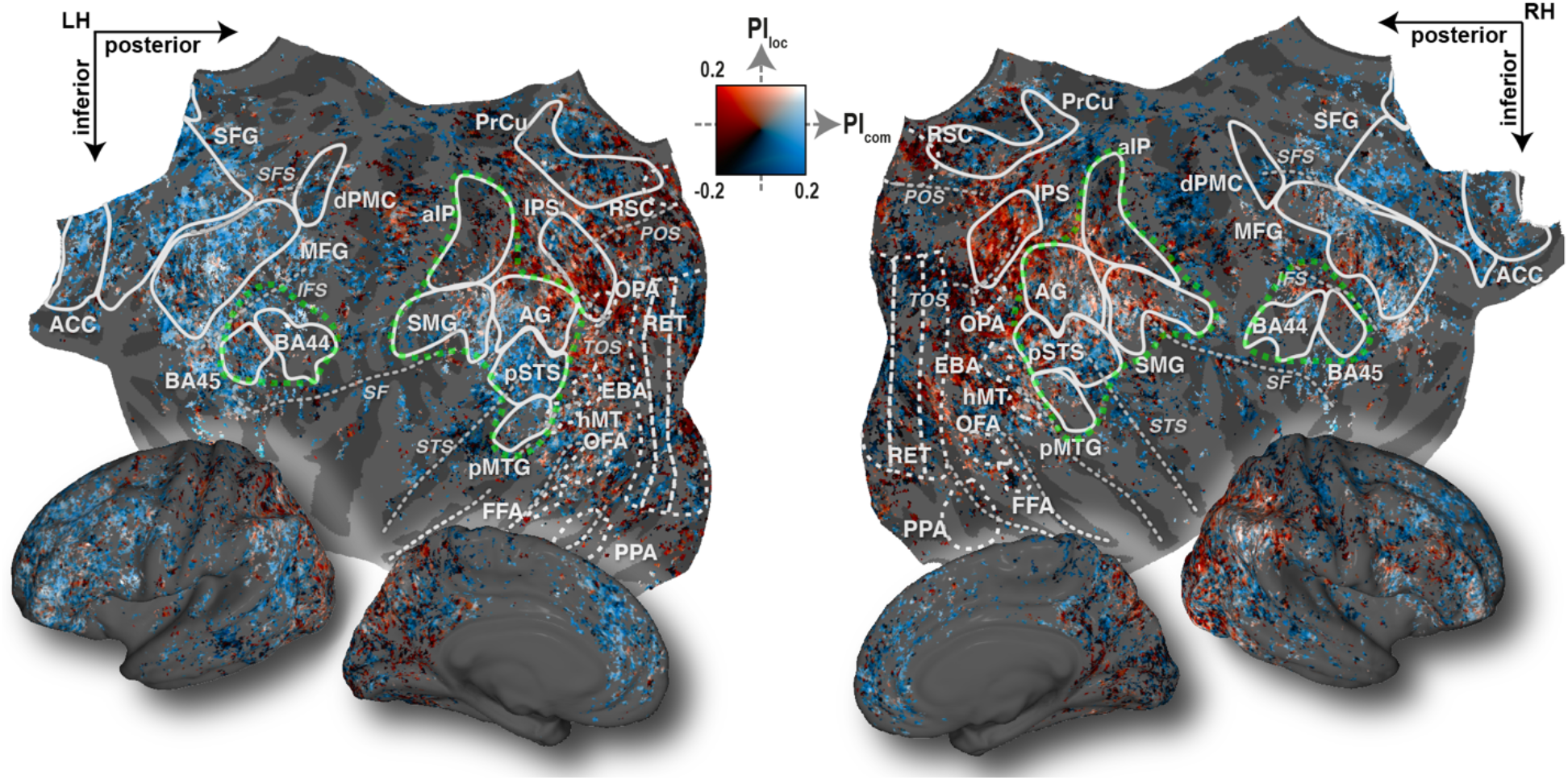
Interaction of tuning shifts with intrinsic selectivity for individual targets. To examine the interaction between tuning shifts and the intrinsic selectivity for individual targets, separate target preference indices (PI) were calculated during search for *communication* (PI_com_), and *locomotion* (PI_loc_) categories. PI during search for a specific target action was taken as the difference in selectivity for the target versus distractor during search for that target. PI_com_ and PI_loc_ values are shown following projection onto the standard brain template. A two-dimensional colourmap was used to annotate each voxel based on PI_com_ and PI_loc_ values (see legend). Figure format is identical to Fig. 3. AON is outlined by green dashed lines. Semantic tuning in voxels across posterior parietal and anterior prefrontal cortices shift toward the attended category irrespective of the search target. However, tuning in many voxels in anterior parietal, occipital, and cingulate cortices shift toward the attended category only during search for communication or only during search for locomotion actions.

**Figure 7.**
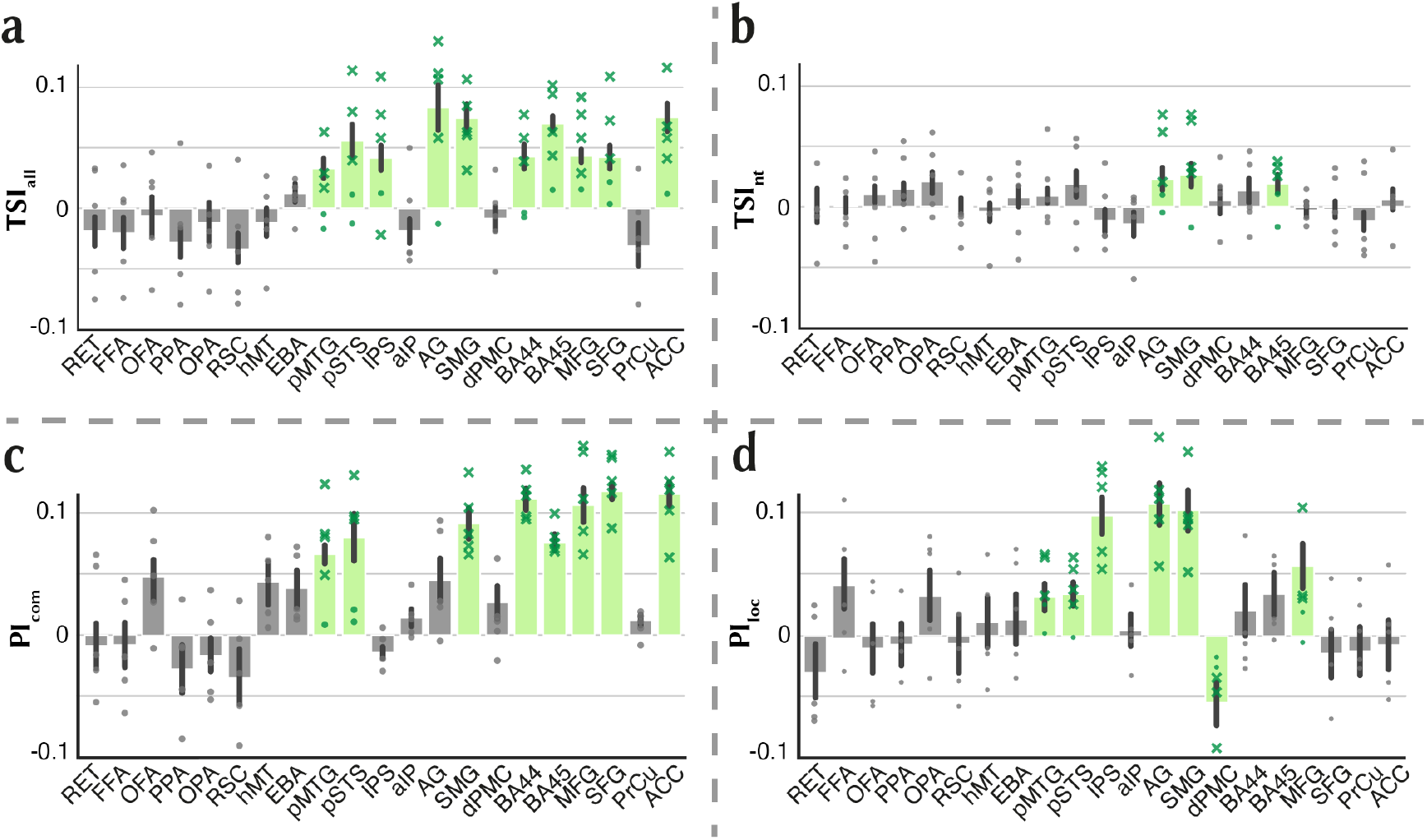
Attentional tuning changes in regions of interest. Average **(a)** TSI_all_, **(b)** TSI_nt_, **(c)** PI_com_, and **(d)** PI_loc_ values were examined in cortical areas (mean±sem across five subjects). Significant values are denoted by green bars and grey bars denote non-significant values (bootstrap test, *p*>0.05). Values for individual subjects are indicated by dots. Grey dots show values in areas with non-significant mean, green dots show non-significant values in areas with significant mean, and green crosses show significant values in areas with significant mean. Tuning shift is significantly greater than zero in many regions across all levels of the AON including occipitotemporal cortex (pSTS, pMTG), posterior parietal cortex (IPS, AG, SMG), and premotor cortex (BA44, BA45), and in regions across prefrontal and cingulate cortices (SFG, ACC). Compared to occipitotemporal areas, attention more diversely modulates semantic representations in parietal and premotor AON nodes, manifested as significantly positive tuning shift for nontarget categories in posterior parietal cortex (AG, SMG) and anterior inferior frontal cortex (BA45). PI_com_ is significantly greater than zero in BA44/45, SFG, and ACC. In contrast, PI_loc_ is significantly greater than zero in IPS and AG and is significantly less than zero in dPMC. Both PI_com_ and PI_loc_ are significantly greater than zero in pSTS, pMTG, SMG, and MFG. Tuning shifts interact with the attention task, and with intrinsic selectivity of cortical areas for target action categories.

#### Areas where both PI_com_ and PI_loc_ are non-significant

We find that PI_com_ and PI_loc_ are non-significant in retinotopic early visual areas (RET; bootstrap test, *p*>0.05) that represent low-level stimulus features, low-level motion-selective area (hMT; *p*>0.05), and object-selective areas (FFA; occipitotemporal face area, OFA; PPA; retrosplenial cortex, RSC; and EBA; *p*>0.05). Furthermore, PI_com_ and PI_loc_ are non-significant in anterior intraparietal cortex (aIP; *p*>0.05), which is involved in representing manipulative actions but not communication or locomotion actions (Noppeney, 2008; Rizzolatti et al., 1997; Urgen and Orban, 2021). These results suggest that during action-based search, semantic tuning shifts least in cortical areas that are selective for lower-level visual features or for neutral high-level action categories irrelevant to the task.

#### Areas where either PI_com_ or PI_loc_ are significant

Several previous studies suggest that lateral and medial prefrontal cortices are causally involved in representing communication actions (Van Overwalle, 2009; Wilson-Mendenhall et al., 2013). Here, we find that PI_com_ is significantly greater than zero in anterior inferior frontal gyrus (BA44/45), in superior frontal gyrus (SFG), and in anterior cingulate cortex (ACC; *p*<0.05). On the other hand, previous reports provide evidence for representation of animate locomotion actions in PPC, including IPS and AG (Abdollahi et al., 2012; Battelli et al., 2003; Bremmer et al., 2001; Ilg et al., 2004). In accord, we find that PI_loc_ is significantly greater than zero in IPS and in AG (*p*<0.05). Taken together, our findings suggest that in areas that are strongly selective for specific action categories, visual search for the preferred action shifts tuning more vigorously towards the preferred target category. It is also worth noting that these attentional effects are not limited to the AON, but rather extend to higher-order cortical areas involved in social cognition. Lastly, we find that PI_loc_ is significantly less than zero while PI_com_ is non-significant (*p*>0.05) in dPMC. This result supports the view that dPMC enhances the representation of distractors during search for locomotion actions (Anticevic et al., 2010; Toepper et al., 2010; Zhou et al., 2012).

#### Areas where both PI_com_ and PI_loc_ are significant

Posterior STS (pSTS), posterior middle temporal gyrus (pMTG), and SMG are considered as AON nodes that maintain representation of actions regardless of their semantic category (Caspers et al., 2010; Jastorff et al., 2016; Lui et al., 2008). We find that both PI_com_ and PI_loc_ are significantly greater than zero in pSTS, pMTG, and SMG (*p*<0.05), consistent with their generic action selectivity. In addition, several previous studies suggest that MFG –as a node in dorsal attention network–facilitates visual search by maintaining the representation of targets (Corbetta and Shulman, 2002; Mars and Grol, 2007; Paneri and Gregoriou, 2017; Ptak et al., 2017). Accordingly, here we find that PI_com_ and PI_loc_ are significantly greater than zero in MFG (*p*<0.05). Overall, these results indicate that in areas with generic action selectivity and in high-level cortical areas, attention facilitates action-based search by shifting representations toward targets irrespective of their semantic category.

The results presented here can be explored online via an interactive brain viewer at http://www.icon.bilkent.edu.tr/brainviewer/shahdloo_etal/.

## Discussion

Here we used brain activity evoked by natural movies to investigate how visual search for action categories modulates semantic representation of a large and diverse set of observed actions across cortex. We examined semantic tuning for 109 action categories in single cortical voxels and quantified their changes as the search target was varied. Importantly, we find that attentional modulation of semantic representations –as manifested in single-voxel semantic tuning shifts– are not restricted to the action observation network, but also extend to higher-level areas in frontoparietal and cingulate cortices. Our results also suggest an interaction between semantic tuning shifts and intrinsic selectivity of neural populations for action categories; semantic representations are most prominently shifted toward a target action in cortical areas that are intrinsically involved in representation of that category. Our results indicate that attention facilitates action perception by modulating semantic tuning for action categories in favour of the targets. These results offer new insights into the effects of category-based visual search on brain responses (Çukur et al., 2013; Erez and Duncan, 2015; Harel et al., 2014; Peelen et al., 2009).

Several previous studies have reported response modulations during action-based attention in parietal and prefrontal cortices, but not in occipitotemporal areas (Nastase et al., 2017, 2018; Nicholson et al., 2017). Yet here we observe significant attentional tuning shifts in occipitotemporal cortex. Unlike previous studies, our analysis approach enables us to measure single-voxel tuning shifts. Moreover, the rich movie stimulus used here contains a large set of action categories that are performed in their natural context, in contrast to the controlled experiments where a handful of actions are displayed on a homogeneous background. Lastly, the set of actions investigated here is restricted to actions performed by animate actors, known to elicit robust responses across the occipitotemporal cortex (Isik et al., 2017; Thompson and Parasuraman, 2012; Walbrin and Koldewyn, 2019; Walbrin et al.,2018). It is thus possible that these design factors have enabled us to detect tuning shifts in early stages of AON comprising occipitotemporal areas.

Recent studies have emphasized the role of AG and SMG in multi-modal semantic representation of actions while observing actions, hearing action sounds, or reading action words (Bedny and Caramazza, 2011; van Dam et al., 2010; Liljeström et al., 2008; Pizzamiglio et al., 2005). There is also evidence suggesting that during semantic processing these areas act as central connectivity hubs, passing the information from low-level perceptual processing areas onto higher-level cortical areas in prefrontal cortex (Farahibozorg et al., 2019; Hoeren et al., 2013). Here, we find significant tuning shifts toward targets in AG and SMG, higher than that in occipitotemporal and premotor AON nodes, irrespective of the search target. Therefore, our results can be taken to suggest a higher place for AG and SMG in the hierarchy of semantic representations of actions compared to remaining AON nodes.

Recent studies suggest that during visual search, cortical areas that are selective for a given object category retain their tuning for the preferred category even when a non-preferred category is the search target (Çukur et al., 2013; Reddy and Kanwisher, 2007; Shahdloo et al., 2020). In this study, by investigating semantic representations during search for individual targets, we find that semantic tuning of voxels in superior parietal cortex –which is suggested to be involved in representation of locomotion actions– are shifted toward locomotion actions only during search for this target. Likewise, semantic tuning of voxels in communication-selective anterior prefrontal cortex are shifted toward communication actions only during search for communication. Taken together, these results suggest that semantic tuning shifts interact with the intrinsic selectivity for target categories.

Here, we derived an embedding space that encodes semantic variability among animate action categories. The first –and most important– dimension in this semantic space distinctively represents self-actions (e.g., chew, yawn) versus actions that involve distal objects or people (e.g., hit, reach). Several recent neuroimaging studies report that observing actions that involve distal objects versus actions without objects lead to non-overlapping population responses in the AON (Handjaras et al., 2015; Tarhan and Konkle, 2020; Wurm and Caramazza, 2019). Specifically, several meta-analyses (Caspers et al., 2010; Grosbras et al., 2012) report that observing actions that involve distal objects leads to increased activation across parietal and prefrontal AON nodes (Buccino et al., 2001; Newman-Norlund et al., 2010) –areas that have been suggested to encode information pertaining to action goals (Grafton and de C Hamilton, 2007; Hamilton and Grafton, 2006; Ramsey and Hamilton, 2010). Thus, our results here can be taken to suggest that distal-proximal distinction is a central feature for representation of action categories in the brain, likely because it helps to disambiguate action goals.

The distal-proximal distinction observed here also converges with an important recent study suggesting that action target is a characteristic feature for action representation (Tarhan and Konkle, 2020). Note, however, that major differences exist between the experimental approaches of the two studies. Tarhan and Konkle compiled a stimulus of video clips depicting a single isolated action, and they have examined brain responses at the level of agent body parts and targets. In contrast, we used cluttered natural movie clips containing multiple actions, actors and targets, and we further used a 109-dimensional feature space to explicitly characterize action category responses. It is nontrivial to pose a model that integrates the effects of multiple agents or targets within cluttered visual scenes, so we cannot directly examine the feature space proposed by Tarhan and Konkle (2020). However, it remains important future work to investigate the contribution of these alternate features to the attentional changes reported here.

Since our measurements are naturally limited by the spatiotemporal resolution of BOLD responses, we cannot make definitive inferences about the neural mechanisms underlying tuning shifts in single voxels. That said, several candidate mechanisms could have contributed to the reported tuning shifts. Previous reports suggest that attention modulates response baseline, response gain, and selectivity in single neurons (Connor et al., 1997; David et al., 2008; Reynolds et al., 2000). Further electrophysiological work would be needed to characterize neural tuning shifts during natural visual search for actions.

The natural movie stimuli used here have greater ecological validity compared to simplified or controlled movie clips used in many previous action-perception studies. That said, action categories in natural movies might be correlated with low-level features such as global motion-energy (Nishimoto et al., 2011; Weiss et al., 2006) and intermediate-level features such as scene dynamics (Grossman and Blake, 2002). If these spurious correlations are substantial, they can confound the estimated action category responses and tuning shifts. We employed several procedures to control for potential biases. First, to minimize correlations between estimated action category responses and global motion-energy of the movie clips, we used a nuisance motion-energy regressor in our modelling procedure (Nishimoto et al., 2011). Second, we restricted analyses to voxels in which the category model explained unique response variance after accounting for low-level motion-energy features and intermediate-level STIP features. However, we do not rule out the possibility that there might be residual influences due to other high-level action features such as expected action goals (Hudson et al., 2016a, 2016b), and actors’ perceived attitude (Bach and Schenke, 2017). Further work is needed to functionally dissociate potential contributions of these high-level features and attentional modulations in action representation.

In conclusion, we showed that natural visual search for a specific action category modulates semantic representations, causing tuning shifts toward the target in single voxels within and beyond the action observation network. Attentional modulations further interact with intrinsic selectivity of neural populations for search targets. This dynamic attentional mechanism can facilitate action perception by efficiently allocating neural resources to accentuate the representation of task-relevant action categories. Overall, our results help explain humans’ astounding ability to perceive others’ actions in dynamic, cluttered daily-life experiences.

## Experimental Procedures

### Subjects

Five healthy adult volunteers with normal or corrected-to-normal vision participated in this study: S1 (male, age 31), S2 (male, age 27), S3 (female, age 32), S4 (male, age 33), S5 (male, age 27). Data were collected at the University of California, Berkeley. The experimental protocol was approved by the Committee for the Protection of Human Subjects at the University of California, Berkeley. All participants gave written informed consent before scanning.

### Stimuli and experimental design

Data for the main experiment were collected in six 10min 50s runs in a single session. Continuous natural movies were used as the stimulus in the main experiment. Three distinct 10min movie segments were compiled from short movie clips (10-20secs) without sound. Movie clips were selected from a diverse set of natural movies (see Nishimoto et al. (2011) for details). Movie clips were cropped into a square frame and downsampled to 512×512px. The movie stimulus was displayed at 15Hz on an MRI-compatible projector screen that covered 24°x24° visual angle. Subjects were instructed to covertly search for target categories in the movies while maintaining fixation. A set of instructions regarding the experimental procedure and exemplars of the search targets were provided to the subjects before the experiment. A colour square of 0.16°x0.16° at the centre with colour changing at 1Hz was used as the fixation spot. A cue word was displayed before each run to indicate the attention target: *communication* or *locomotion*. The *communication* target contained actions with the intent of communication, including both verbal communication actions and nonverbal gestural communication actions (e.g., talking, shouting, smirking). The *locomotion* target contained locomotion-related actions with the intent of moving animate entities, including humans and anthropomorphized animals (e.g., moving, running, driving). The order of attention conditions was interleaved across runs to minimize subject expectation bias. This resulted in presentation of 1800sec of movies without repetition in each attention condition. Data from the first 20secs and last 30secs of each run were discarded to minimize effects of transient confounds. Following these procedures, 900 data samples for each attention condition were obtained.

A separate set of functional data were collected while subjects passively viewed 120min of natural movies (i.e., passive-viewing data; Huth et al., 2012). This dataset was used to construct the semantic space and to select voxels subjected to further analyses. Data for the passive-viewing experiment were collected in twelve 10min 50s runs in which 12 separate movie segments were displayed. Presentation procedures were the same between the main experiment and passive-viewing experiment, save for the number of runs. The passive-viewing dataset contained 3600 data samples.

### Category features

A category feature space was constructed to encode the information pertaining to object and action categories in the movies. Each second of the movie stimulus was manually labelled using the WordNet lexicon (Miller, 1995) to find the time course for presence of 922 different object and action categories in the movie stimulus. This yielded an indicator matrix where each row represents a one-second clip of the movie stimulus and each column represents a category. Finally, category features were obtained by downsampling the indicator matrix to 0.5Hz in order to match the acquisition rate of fMRI.

### Motion-energy features

To infer cortical selectivity for low-level scene features, local spatial frequency and orientation information of each frame of the movie stimulus were quantified using a motion-energy filter bank. The filter bank contained 2139 Gabor filters that were computed at eight directions (0 to 350°, in 45° steps), three temporal frequencies (0, 2, and 4Hz), and six spatial frequencies (0, 1.5, 3, 6, 12, and 24 cycles/image). Filters were placed on a square grid spanning the 24°x24° field of view. The luminance channel was extracted from the movie frames and passed through the filter bank. The outputs were then passed through a compressive nonlinearity to yield the motion-energy features (Lescroart and Gallant, 2019; Nishimoto et al., 2011). Finally, the motion-energy features were temporally downsampled to match the fMRI acquisition rate.

### Space-time Interest Points (STIP) features

Intermediate-level kinematic information of the movies were quantified by constructing the Space-Time Interest Point (STIP) features using STIP toolbox (Laptev, 2005; Laptev et al., 2008). STIP features have been successfully leveraged in many computer vision applications to recognize human actions. As detailed in Laptev (2005) and Laptev et al. (2008), Harris operators (Harris and Stephens, 1988) were used to identify spatiotemporal interest points in the movie stimulus at multiple scales 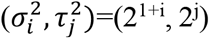, *i* ∈ {1,…,6}, *j* ∈ {1,2}, where *σ* and *τ* are the standard deviations of the Gaussian kernels in spatial and temporal domains respectively. Histograms of oriented gradients (HoG; Dalal and Triggs, 2005), and histograms of optical flow (HoF; Holte et al., 2010) were calculated in the (*Δ_x,i_*Δ_*y,i*_,Δ_*t,j*_) spatiotemporal neighbourhood of each interest point, where *Δ_x,i_* = Δ_*y,i*_ = 2*kσ_i_* and *Δ_t,j_* = 2*kτ_j_*, and *k* is the scale factor. Scale factor was set to 9 according to the default configuration of the toolbox. Finally, normalized histograms were concatenated to construct the collection of 162 STIP features and were downsampled to match the acquisition rate of fMRI.

### Model estimation and testing

For each voxel, separate linearized models were estimated to relate each feature representation to the BOLD responses as detailed in *Supplementary Methods* and in previous studies from our lab (Çukur et al., 2013; Kiremitçi et al., 2021; Shahdloo et al., 2020). Specifically, category models were fit to estimate category response vectors that represented the contribution of each category to single-voxel BOLD responses. Furthermore, a motion energy model and a STIP model were fit in each voxel to represent the contribution of the low- and intermediate-level stimulus features to the responses. These alternative models were further used to select analysis voxels (i.e., semantic voxels), as detailed in *Supplementary Methods*.

### Action category responses

The fit category responses reflect voxel tuning for each of the 922 object and action categories in the movie stimulus. To infer tuning for action categories, 922-dimensional category responses were masked to select only the 109 action categories. This yielded the voxelwise 109-dimensional action category responses.

### Semantic representation of actions

Passive-viewing data were used to construct a continuous semantic space for action category representation. In this space, semantically similar action categories would project to nearby points, whereas semantically dissimilar categories would project to distant points (Huth et al., 2012). Category models were fit and action category responses during passive viewing were estimated. A group semantic space was then obtained using principal component analysis (PCA) on the action category responses of cortical voxels pooled across all subjects. To maximize the quality of the semantic space, voxels in which the category model predicted unique response variance after accounting for the variance attributed to low- and intermediate-level stimulus features were selected. These voxels were further refined to include only the top 3,000 best predicted voxels within each subject. The top 12 principal components (PCs) that explained more than 95% of the variance in responses were selected. Subsequent analyses were also repeated using the top 8 PCs that explained more than 90% of the response variance but the results remained consistent. Semantic tuning profile for each voxel under each search task was then obtained by projecting the respective action category responses onto the PCs. To facilitate the visualisation of the semantic space, action categories were clustered (see Fig. 4 and *Supplementary Methods*).

### Consistency of the semantic space across subjects

To test whether the estimated semantic space is consistent across subjects, we used a leave-one-out cross-validation procedure. In each cross-validation fold, voxels from four subjects were used to derive 12 PCs to construct a semantic space. In the left-out subject, semantic tuning profile for each voxel was obtained by projecting action category responses during passive viewing onto the derived PCs. Next, product-moment correlation coefficient was calculated between the tuning profiles in the derived space and the tuning profiles in the original semantic space. Results were averaged across semantic voxels in the left-out subject. The cross-validated semantic spaces consistently correlate with the original semantic space (Supp. Fig. 3).

### Characterizing tuning shifts

Attentional tuning shifts toward or away from targets would be reflected in modulation of semantic selectivity for *communication* or *locomotion* action categories. Thus, the magnitude and direction of tuning shifts can be assessed by comparing the semantic selectivity for these categories between the two search tasks. Semantic selectivity for the two target categories was quantified as the similarity between semantic tuning profiles and idealized templates tuned solely for *communication* or *locomotion* action categories. First, idealized category responses were constructed as 109-dimensional vectors that contained ones for target categories (either *communication* or *locomotion* categories) and zeros elsewhere. Idealized templates were then obtained by projecting these idealized category responses onto the semantic space. Semantic selectivity for each target category was quantified as product-moment correlation coefficient between voxel semantic tuning profile and the corresponding template

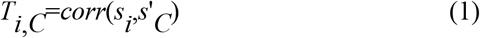

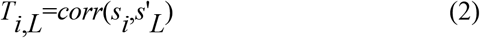

where T_i,C_ and T_i,L_ are the tuning strength for *communication* and *locomotion* during condition *i*∈{C, L} denoting attend to *communication* or attend to *locomotion*, s_i_ is the semantic tuning profile during condition *i*, and s’_C_ and s’_L_ denote the idealized semantic tuning templates for *communication* and *locomotion*, respectively. Finally, voxelwise tuning shift index (TSI_all_) was quantified as

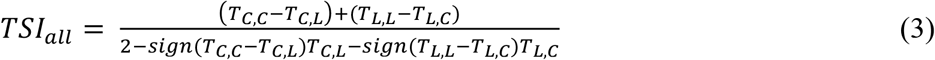

Tuning shifts toward the attended category would yield positive values where a TSI_all_ of 1 indicates a complete match between voxel semantic tuning and idealized templates, whereas negative values would indicate shifts away from the attended category where a TSI_all_ of −1 indicates a complete mismatch between voxel tuning and idealized templates. A TSI_all_ of 0 would indicate that the voxel tuning did not shift between the two search tasks. To investigate attentional modulation of semantic tuning for nontarget categories, a separate tuning shift index was calculated (TSI_nt_). To calculate TSI_nt_, action category responses were masked to select the nontarget categories (i.e., all actions except communication and locomotion categories) prior to projection onto the semantic space. To study the tuning shifts in an ROI, TSIs were averaged across semantic voxels within the ROI.

### Characterizing target preference during visual search

To investigate the interaction between tuning shifts and intrinsic selectivity for individual target action categories, we quantified a target preference index (PI∈[-1,1]) separately during search for communication actions (PI_com_) and during search for locomotion actions (PI_loc_). PI during search for each target action was taken as the difference in selectivity for the attended versus the unattended target

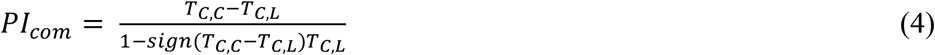

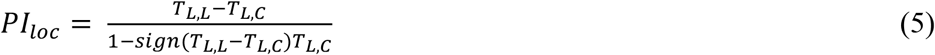

where PI_com_ denotes the relative tuning preference for communication actions during search for communication, and PI_loc_ denotes the relative tuning preference for locomotion actions during search for locomotion. In this scheme, a PI of 1 indicates a complete match between voxel semantic tuning and the idealized template for the target, whereas a PI of −1 indicates a complete mismatch between voxel tuning and the idealized template for the target. Finally, a PI of 0 indicates that the voxel semantic tuning does not shift toward any of the target actions.

## Data and software availability

Data supporting the findings of this study are available from the corresponding authors upon request. Results can be explored online via an interactive brain viewer at http://www.icon.bilkent.edu.tr/brainviewer/shahdloo_etal/.

The codes used to estimate spatially informed voxelwise model weights is freely available on GitHub at https://github.com/icon-lab/SPIN-VM.

## Author contributions

Conceptualization, J.L.G and T.C.; Methodology, M.S., and T.C.; Software, M.S., and E.C.; Investigation, M.S., B.A.U. and T.C.; Writing-Original Draft, M.S.; Writing-Review and Editing, M.S., T.C., B.A.U., J.L.G, and E.C.; Funding Acquisition, T.C.; Resources, M.S., T.C., and J.L.G; Supervision, T.C.

The authors declare no competing interests.

## References

Abdollahi, R.O., Jastorff, J., and Orban, G.A. (2012). Common and Segregated Processing of Observed Actions in Human SPL. Cerebral Cortex 23, 2734–2753.

Anticevic, A., Repovs, G., and Barch, D.M. (2010). Resisting emotional interference: Brain regions facilitating working memory performance during negative distraction. Cognitive, Affective, & Behavioral Neuroscience 10, 159–173.

Bach, P., and Schenke, K.C. (2017). Predictive social perception: Towards a unifying framework from action observation to person knowledge. Social and Personality Psychology Compass 11, e12312.

Battelli, L., Cavanagh, P., and Thornton, I.M. (2003). Perception of biological motion in parietal patients. Neuropsychologia 41, 1808–1816.

Bedny, M., and Caramazza, A. (2011). Perception, action, and word meanings in the human brain: The case from action verbs. Annals of the New York Academy of Sciences 1224, 81–95.

Bremmer, F., Schlack, A., Duhamel, J.-R., Graf, W., and Fink, G.R. (2001). Space Coding in Primate Posterior Parietal Cortex. NeuroImage 14, S46–S51.

Buccino, G., Binkofski, F., Fink, G.R., Fadiga, L., Fogassi, L., Gallese, V., Seitz, R.J., Zilles, K., Rizzolatti, G., and Freund, H.J. (2001). Action observation activates premotor and parietal areas in a somatotopic manner: An fMRI study. European Journal of Neuroscience 13, 400–404.

Caspers, S., Zilles, K., Laird, A.R., and Eickhoff, S.B. (2010). ALE meta-analysis of action observation and imitation in the human brain. NeuroImage 50, 1148–1167.

Cavina-Pratesi, C., Connolly, J.D., Monaco, S., Figley, T.D., Milner, D., Schenk, T., Culham, J.C. (2018). Human neuroimaging reveals the subcomponents of grasping, reaching and pointing actions. Cortex 98, 128–148.

Çelik, E., Dar, S.U.H., Yılmaz, Ö., Keleş, Ü., and Çukur, T. (2019). Spatially informed voxelwise modeling for naturalistic fMRI experiments. NeuroImage 186, 741–757.

Connor, C.E., Preddie, D.C., Gallant, J.L., and Van Essen, D.C. (1997). Spatial Attention Effects in Macaque Area V4. Journal of Neuroscience 17, 3201–3214.

Corbetta, M., and Shulman, G.L. (2002). Control of goal-directed and stimulus-driven attention in the brain. Nature Reviews Neuroscience 3, 215–229.

Corbo, D., and Orban, G.A. (2017). Observing others speak or sing activates SPT and neighboring parietal cortex. Journal of Cognitive Neuroscience 29, 1002–1021.

Çukur, T., Nishimoto, S., Huth, A.G., and Gallant, J.L. (2013). Attention during natural vision warps semantic representation across the human brain. Nature Neuroscience 16, 763–770.

Dalal, N., and Triggs, B. (2005). Histograms of oriented gradients for human detection. In IEEE Computer Society Conference on Computer Vision and Pattern Recognition (CVPR’05), IEEE, 886–893.

van Dam, W.O., Rueschemeyer, S.-A., and Bekkering, H. (2010). How specifically are action verbs represented in the neural motor system: An fMRI study. NeuroImage 53, 1318–1325.

David, S.V., Hayden, B.Y., Mazer, J.A., and Gallant, J.L. (2008). Attention to stimulus features shifts spectral tuning of V4 neurons during natural vision. Neuron 59, 509–521.

Erez, Y., and Duncan, J. (2015). Discrimination of Visual Categories Based on Behavioral Relevance in Widespread Regions of Frontoparietal Cortex. Journal of Neuroscience 35, 12383–12393.

Farahibozorg, S.-R., Henson, R.N., Woollams, A.M., and Hauk, O. (2019). Distinct roles for the Anterior Temporal Lobe and Angular Gyrus in the spatio-temporal cortical semantic network. BioRxiv, 544114.

Ferri, S., Rizzolatti, G., and Orban, G.A. (2015). The organisation of the posterior parietal cortex devoted to upper limb actions: An fMRI study. Human Brain Mapping 36, 3845–3866.

Grafton, S.T., and de C Hamilton, A.F. (2007). Evidence for a distributed hierarchy of action representation in the brain. Human Movement Science 26, 590–616.

Grosbras, M.H., Beaton, S., and Eickhoff, S.B. (2012). Brain regions involved in human movement perception: A quantitative voxel-based meta-analysis. Human Brain Mapping 33, 431–454.

Grossman, E.D., and Blake, R. (2002). Brain areas active during visual perception of biological motion. Neuron 35, 1167–1175.

Hamilton, A.F. de C., and Grafton, S.T. (2006). Goal representation in human anterior intraparietal sulcus. Journal of Neuroscience 26, 1133–1137.

Handjaras, G., Bernardi, G., Benuzzi, F., Nichelli, P.F., Pietrini, P., and Ricciardi, E. (2015). A topographical organisation for action representation in the human brain. Human Brain Mapping 36, 3832–3844.

Harel, A., Kravitz, D.J., and Baker, C.I. (2014). Task context impacts visual object processing differentially across the cortex. PNAS 111, E962–71.

Harris, C., and Stephens, M. (1988). A combined corner and edge detector. In In Proc. Fourth Alvey Vision Conference, 147–152.

Haxby, J.V., Connolly, A.C., and Guntupalli, J.S. (2014). Decoding neural representational spaces using multivariate pattern analysis. Annual Reviews of Neuroscience 37, 435–456.

Herrington, J., Nymberg, C., Faja, S., Price, E., and Schultz, R. (2012). The responsiveness of biological motion processing areas to selective attention towards goals. NeuroImage 63, 581–590.

Hoeren, M., Kaller, C.P., Glauche, V., Vry, M.-S., Rijntjes, M., Hamzei, F., and Weiller, C. (2013). Action semantics and movement characteristics engage distinct processing streams during the observation of tool use. Experimental Brain Research 229, 243–260.

Holte, M.B., Moeslund, T.B., and Fihl, P. (2010). View-invariant gesture recognition using 3D optical flow and harmonic motion context. Computer Vision and Image Understanding 114, 1353–1361.

Hudson, M., Nicholson, T., Ellis, R., and Bach, P. (2016a). I see what you say: Prior knowledge of other’s goals automatically biases the perception of their actions. Cognition 146, 245–250.

Hudson, M., Nicholson, T., Simpson, W.A., Ellis, R., and Bach, P. (2016b). One step ahead: The perceived kinematics of others’ actions are biased toward expected goals. Journal of Experimental Psychology: General 145, 1–7.

Huth, A.G., Nishimoto, S., Vu, A.T., and Gallant, J.L. (2012). A Continuous Semantic Space Describes the Representation of Thousands of Object and Action Categories across the Human Brain. Neuron 76, 1210–1224.

Ilg, U.J., Schumann, S., and Thier, P. (2004). Posterior parietal cortex neurons encode target motion in world-centered coordinates. Neuron 43, 145–151.

Isik, L., Koldewyn, K., Beeler, D., and Kanwisher, N.G. (2017). Perceiving social interactions in the posterior superior temporal sulcus. PNAS 114, E9145–E9152.

Jastorff, J., and Orban, G.A. (2009). Human Functional Magnetic Resonance Imaging Reveals Separation and Integration of Shape and Motion Cues in Biological Motion Processing. Journal of Neuroscience 29, 7315–7329.

Jastorff, J., Begliomini, C., Fabbri-Destro, M., Rizzolatti, G., and Orban, G.A. (2010). Coding observed motor acts: Different organisational principles in the parietal and premotor cortex of humans. Journal of Neurophysiology 104, 128–140.

Jastorff, J., Abdollahi, R.O., Fasano, F., and Orban, G.A. (2016). Seeing biological actions in 3D: An fMRI study. Human Brain Mapping 37, 203–219.

Johansson, G. (1973). Visual perception of biological motion and a model for its analysis. Perception & Psychophysics 14, 201–211.

Kilintari, M., Raos, V., and Savaki, H.E. (2014). Involvement of the Superior Temporal Cortex in Action Execution and Action Observation. Journal of Neuroscience 34, 8999–9011.

Kiremitçi, I., Yilmaz, Ö., Çelik, E., Shahdloo, M., Huth, A.G., and Çukur, T. (2021). Attentional Modulation of Hierarchical Speech Representations in a Multi-Talker Environment. Cerebral Cortex, bhab136

Kriegeskorte, N., and Kievit, R.A. (2013). Representational geometry: Integrating cognition, computation, and the brain. Trends in Cognitive Sciences 17, 401–412.

Kriegeskorte, N., Mur, M., and Bandettini, P.A. (2008). Representational similarity analysis - connecting the branches of systems neuroscience. Frontiers in Systems Neuroscience 2, 4.

de Lange, F.P., Spronk, M., Willems, R.M., Toni, I., and Bekkering, H. (2008). Complementary Systems for Understanding Action Intentions. Current Biology 18, 454–457.

Laptev, I. (2005). On space-time interest points. International Journal of Computer Vision, 107–123.

Laptev, I., Marszałek, M., Schmid, C., and Rozenfeld, B. (2008). Learning realistic human actions from movies. 26th IEEE Conference on Computer Vision and Pattern Recognition (CVPR), 1–8.

Lescroart, M.D., and Gallant, J.L. (2019). Human Scene-Selective Areas Represent 3D Configurations of Surfaces. Neuron 101, 178–192.e7.

Liljeström, M., Tarkiainen, A., Parviainen, T., Kujala, J., Numminen, J., Hiltunen, J., Laine, M., and Salmelin, R. (2008). Perceiving and naming actions and objects. NeuroImage 41, 1132–1141.

Lui, F., Buccino, G., Duzzi, D., Benuzzi, F., Crisi, G., Baraldi, P., Nichelli, P., Porro, C.A., and Rizzolatti, G. (2008). Neural substrates for observing and imagining non-object-directed actions. Social Neuroscience 3, 261–275.

Mars, R.B., and Grol, M.J. (2007). Dorsolateral prefrontal cortex, working memory, and prospective coding for action. Journal of Neuroscience 27, 1801–1802.

Miller, G.A. (1995). WordNet: a lexical database for English. Communications of ACM 38, 39–41.

Molinari, E., Baraldi, P., Campanella, M., Duzzi, D., Nocetti, L., Pagnoni, G., and Porro, C.A. (2013). Human parietofrontal networks related to action observation detected at rest. Cerebral Cortex 23, 178–186.

Muthukumaraswamy, S.D., and Singh, K.D. (2008). Modulation of the human mirror neuron system during cognitive activity. Psychophysiology 45, 896–905.

Muthukumaraswamy, S.D., Johnson, B.W., and McNair, N.A. (2004). Mu rhythm modulation during observation of an object-directed grasp. Cognitive Brain Research 19, 195–201.

Nastase, S.A., Connolly, A.C., Oosterhof, N.N., Halchenko, Y.O., Guntupalli, J.S., Di Oleggio Castello, M.V., Gors, J., Gobbini, M.I., and Haxby, J.V. (2017). Attention selectively reshapes the geometry of distributed semantic representation. Cerebral Cortex 27, 4277–4291.

Nastase, S.A., Halchenko, Y.O., Connolly, A.C., Gobbini, M.I., and Haxby, J.V. (2018). Neural responses to naturalistic clips of behaving animals in two different task contexts. Frontiers in Neuroscience 12, 316.

Nelissen, K., Vanduffel, W., and Orban, G.A. (2006). Charting the Lower Superior Temporal Region, a New Motion-Sensitive Region in Monkey Superior Temporal Sulcus. Journal of Neuroscience 26, 5929–5947.

Nelissen, K., Borra, E., Gerbella, M., Rozzi, S., Luppino, G., Vanduffel, W., Rizzolatti, G., and Orban, G.A. (2011). Action observation circuits in the macaque monkey cortex. Journal of Neuroscience 31, 3743–3756.

Newman-Norlund, R., van Schie, H.T., van Hoek, M.E.C., Cuijpers, R.H., and Bekkering, H. (2010). The role of inferior frontal and parietal areas in differentiating meaningful and meaningless object-directed actions. Brain Research 1315, 63–74.

Nicholson, T., Roser, M., and Bach, P. (2017). Understanding the Goals of Everyday Instrumental Actions Is Primarily Linked to Object, Not Motor-Kinematic, Information: Evidence from fMRI. PLoS ONE 12, e0169700.

Nili, H., Cai W., Alexander W., Li S., William M. W., and Kriegeskorte N. (2014). A Toolbox for Representational Similarity Analysis. PLoS Computational Biology 10, e1003553.

Nishimoto, S., Vu, A.T., Naselaris, T., Benjamini, Y., Yu, B., and Gallant, J.L. (2011). Reconstructing Visual Experiences from Brain Activity Evoked by Natural Movies. Current Biology 21, 1641–1646.

Noppeney, U. (2008). The neural systems of tool and action semantics: A perspective from functional imaging. Journal of Physiology 102, 40–49.

Oberman, L.M., Pineda, J.A., and Ramachandran, V.S. (2007). The human mirror neuron system: A link between action observation and social skills. Social Cognitive and Affective Neuroscience 2, 62–66.

Oosterhof, N.N., Wiggett, A.J., Diedrichsen, J., Tipper, S.P., and Downing, P.E. (2010). Surface-based information mapping reveals crossmodal vision-action representations in human parietal and occipitotemporal cortex. Journal of Neurophysiology 104, 1077–1089.

Oosterhof, N.N., Tipper, S.P., and Downing, P.E. (2012). Viewpoint (in)dependence of action representations: an MVPA study. Journal of Cognitive Neuroscience 24, 975–989.

Oosterhof, N.N., Tipper, S.P., and Downing, P.E. (2013). Crossmodal and action-specific: Neuroimaging the human mirror neuron system. Trends in Cognitive Sciences 17, 311–318.

Paneri, S., and Gregoriou, G.G. (2017). Top-down control of visual attention by the prefrontal cortex. Functional specialization and long-range interactions. Frontiers in Neuroscience 11, 545.

Peelen, M.V., Fei-Fei, L., and Kastner, S. (2009). Neural mechanisms of rapid natural scene categorization in human visual cortex. Nature 460, 94–97.

Pizzamiglio, L., Aprile, T., Spitoni, G., Pitzalis, S., Bates, E., D’Amico, S., and Di Russo, F. (2005). Separate neural systems for processing action- or non-action-related sounds. NeuroImage 24, 852–861.

Ptak, R., Schnider, A., and Fellrath, J. (2017). The Dorsal Frontoparietal Network: A Core System for Emulated Action. Trends in Cognitive Sciences 21, 589–599.

Puglisi, G., Leonetti, A., Landau, A., Fornia, L., Cerri, G., and Borroni, P. (2017). The role of attention in human motor resonance. PLoS ONE 12, e0177457.

Puglisi, G., Leonetti, A., Cerri, G., and Borroni, P. (2018). Attention and cognitive load modulate motor resonance during action observation. Brain & Cognition 128, 7–16.

Ramsey, R., and Hamilton, A.F.D.C. (2010). Understanding actors and object-goals in the human brain. NeuroImage 50, 1142–1147.

Reddy, L., and Kanwisher, N.G. (2007). Category Selectivity in the Ventral Visual Pathway Confers Robustness to Clutter and Diverted Attention. Current Biology 17, 2067–2072.

Reynolds, J.H., Pasternak, T., and Desimone, R. (2000). Attention Increases Sensitivity of V4 Neurons. Neuron 26, 703–714.

Rizzolatti, G., and Matelli, M. (2003). Two different streams form the dorsal visual system: Anatomy and functions. Experimental Brain Research 153, 146–157.

Rizzolatti, G., Fogassi, L., and Gallese, V. (1997). Parietal cortex: from sight to action. Current Opinion in Neurobiology 7, 562–567.

Rowe, J., Friston, K., Frackowiak, R., and Passingham, R. (2002). Attention to action: Specific modulation of corticocortical interactions in humans. NeuroImage 17, 988–998.

Rozzi, S., and Fogassi, L. (2017). Neural coding for action execution and action observation in the prefrontal cortex and its role in the organisation of socially driven behavior. Frontiers in Neuroscience 11, 276.

Safford, A.S., Hussey, E.A., Parasuraman, R., and Thompson, J.C. (2010). Object-based attentional modulation of biological motion processing: spatiotemporal dynamics using functional magnetic resonance imaging and electroencephalography. Journal of Neuroscience 30, 9064–9073.

Schuch, S., Bayliss, A.P., Klein, C., and Tipper, S.P. (2010). Attention modulates motor system activation during action observation: Evidence for inhibitory rebound. Experimental Brain Research 205, 235–249.

Seidl, K.N., Peelen, M.V., and Kastner, S. (2012). Neural evidence for distracter suppression during visual search in real-world scenes. Journal of Neuroscience 32, 11812–11819.

Shahdloo, M., Çelik, E., and Çukur, T. (2020). Biased competition in semantic representation during natural visual search. NeuroImage 216, 116383.

Tarhan, L., and Konkle, T. (2020). Sociality and interaction envelope organise visual action representations. Nature Communications 11, 3002.

Thompson, J., and Parasuraman, R. (2012). Attention, biological motion, and action recognition. NeuroImage 59, 4–13.

Toepper, M., Gebhardt, H., Beblo, T., Thomas, C., Driessen, M., Bischoff, M., Blecker, C.R., Vaitl, D., and Sammer, G. (2010). Functional correlates of distractor suppression during spatial working memory encoding. Neuroscience 165, 1244–1253.

Urgen, B.A., and Orban, G.A. (2021). The unique role of parietal cortex in action observation: Functional organisation for communicative and manipulative actions. NeuroImage 273, 118220.

Urgen, B.A., Pehlivan, S., and Saygin, A.P. (2019). Distinct representations in occipito-temporal, parietal, and premotor cortex during action perception revealed by fMRI and computational modeling. Neuropsychologia 127, 35–47.

Van Overwalle, F. (2009). Social cognition and the brain: A meta-analysis. Human Brain Mapping 30, 829–858.

Walbrin, J., and Koldewyn, K. (2019). Dyadic interaction processing in the posterior temporal cortex. NeuroImage 198, 296–302.

Walbrin, J., Downing, P., and Koldewyn, K. (2018). Neural responses to visually observed social interactions. Neuropsychologia 112, 31–39.

Weiss, P.H., Rahbari, N.N., Lux, S., Pietrzyk, U., Noth, J., and Fink, G.R. (2006). Processing the spatial configuration of complex actions involves right posterior parietal cortex: An fMRI study with clinical implications. Human Brain Mapping 27, 1004–1014.

Wilson-Mendenhall, C.D., Simmons, W.K., Martin, A., and Barsalou, L.W. (2013). Contextual processing of abstract concepts reveals neural representations of nonlinguistic semantic content. Journal of Cognitive Neuroscience 25, 920–935.

Wurm, M.F., and Caramazza, A. (2019). Lateral occipitotemporal cortex encodes perceptual components of social actions rather than abstract representations of sociality. NeuroImage 202, 116153.

Wurm, M.F., Caramazza, A., and Lingnau, A. (2017). Action Categories in Lateral Occipitotemporal Cortex Are Organised Along Sociality and Transitivity. Journal of Neuroscience 37, 562–575.

Zhou, X., Katsuki, F., Qi, X.-L., and Constantinidis, C. (2012). Neurons with inverted tuning during the delay periods of working memory tasks in the dorsal prefrontal and posterior parietal cortex. J Neurophysiology 108, 31–38.

