## Supplementary Materials for "Task-Dependent Warping of Semantic Representations During Search for Visual Action Categories"

or

Tolga Çukur, **

### Supplementary Methods

#### List of action categories

smile, communicate, nod, indicate, grimace, read, cell phone, talk, write, argue, shout, pray, gesticulate, yell, smirk, fly, climb, ride horseback, chase, hike, ride, crash, drive, scurry, dive, swim, descend, walk, rise, travel, edge, rush, gallop, pounce, pass, fall, canter, ski, skid, run, crawl, chew, change, grow, lean, arise, break, bend, crouch, struggle, breathe, yawn, huff, sigh, punch, drive, lick, snog, strike, touch, caress, slam, hit, eat, drink, consume, decant, put, kick, lift, push, propel, stand, jab, turn, drag, slide, pull, clap, move, pour, fasten, connect, act, affect, change, stretch, shape, raise, dance, turn, tumble, exit, reach, spin, revolve, jump, flip, hop, bounce, sneak

#### fMRI data collection

Data were collected on a 3T Siemens Tim Trio MRI scanner (Siemens Medical Solutions) via a 32-channel receiver coil. Functional data were collected using a T2*-weighted gradient-echo echo-planar-imaging pulse sequence with the following parameters: TR=2sec, TE=33msec, water-excitation pulse with flip angle=70º, voxel size=2.24mm×2.24mm×4.13mm, field of view=224mm×224mm, 32 axial slices. To construct cortical surfaces, anatomical data were collected using a three-dimensional T1-weighted magnetization-prepared rapid-acquisition gradient-echo (MPRAGE) sequence with the following parameters: TR=2.3sec, TE=3.45 msec, flip angle=10º, voxel size=1mm×1mm×1mm, field of view=256mm×212mm×256mm. Surface flattening and visualisation were done via Freesurfer and PyCortex (Dale et al., 1999; Reuter et al., 2012; Gao et al., 2015).

#### fMRI data preprocessing

Motion correction was performed using Statistical Parametric Mapping toolbox (SPM12; Friston et al., 1995). Functional volumes were aligned to the first image from the first run in each subject. Brain tissue was identified using the brain extraction tool (BET) from the FSL software package (Smith, 2002). Low-frequency response components were detected using a third order Savitzky-Golay low-pass filter with 240sec temporal window and were removed from voxel responses. Voxel responses were then z-scored to attain zero mean and unit variance. Voxels within the 2mm neighbourhood of the cortical sheet were identified as cortical voxels in each subject (S1, 37791 voxels; S2, 32671 voxels; S3, 36942 voxels; S4, 42090 voxels; S5, 39254 voxels).

#### Definition of regions of interest

To define the anatomical regions of interest (ROIs) in each subject, the cortical surface was segmented into 156 regions of the Destrieux atlas (Destrieux et al., 2010) via Freesurfer. Segmentation results were projected from the anatomical space onto the functional space using PyCortex, and each voxel was assigned an anatomical label based on the projections. Functional ROIs were identified in each subject using visual category and retinotopic localizers (Huth et al., 2012). Localizer experiments for visual category-selective areas (fusiform face area, FFA; occipital face area, OFA; parahippocampal place area, PPA; retrosplenial cortex, RSC) were performed in six 4.5 min runs of 16 blocks (Huth et al., 2012). Subjects passively viewed 20 random static images from one of the objects, scenes, body parts, faces, or spatially scrambled objects groups in each block. Each image was shown for 300ms following a 500ms blank period. PPA and RSC were identified as voxels with positive scene versus objects contrast (*t*-test, $p<10^{-4}$, uncorrected). FFA and OFA were defined using face-versus-object contrast (*t*-test, $p<10^{-4}$, uncorrected). The boundaries of these areas were hand drawn on the cortical surfaces along the contours at which the contrast level reached half of the maximum. Localizer experiment for early visual areas (RET: V1, V2, V3) contained four 9min runs. Subjects viewed clockwise and counter-clockwise rotating polar wedges in two runs. In the remaining two runs, subjects viewed expanding and contracting rings. Visual angle and eccentricity maps were used to define visual areas V1-3. Finally, ROIs were refined to voxels inside the drawn boundaries near a 2mm neighbourhood of the cortical sheet.

#### Abbreviations for regions of interest and important sulci

Several regions of interest and important sulci were labelled on the flattened cortical surfaces to guide the reader.

***Regions of interest*:** pMTG, posterior middle temporal gyrus; pSTS, posterior superior temporal sulcus; AG, angular gyrus; SMG, supramarginal gyrus; IPS, intraparietal sulcus; aIP, anterior intraparietal cortex; PrCu, precuneous; dPMC, dorsal premotor cortex; BA44/45, Brodmann area 44/45; MFG, middle frontal gyrus; SFG, superior frontal gyrus; ACC, anterior cingulate cortex; RET, early visual areas V1-3; FFA, fusiform face area; OFA, occipital face area; PPA, parahippocampal place area; RSC, retrosplenial cortex.

***Sulci*:** TOS, temporo-occipital sulcus; STS, superior temporal sulcus; SF, Sylvian fissure; IFS, inferior frontal sulcus; MFS, middle frontal sulcus; SFS, superior frontal sulcus.

#### Head motion, eye-movement, and physiological noise

To prevent head motion and physiological noise confounds, estimates of these nuisance factors were regressed out of the BOLD responses. Six affine motion time courses estimated during the motion-correction stage were taken as the head-motion regressors. The cardiac and respiratory activity during the main experiment were recorded using a pulse oximeter and a pneumatic belt. These data were then used to estimate two regressors to capture respiration and nine regressors to capture cardiac activity (Verstynen and Deshpande, 2011).

To ensure that eye-movements did not unduly bias the results, several control analyses were performed. ViewPoint EyeTracker (Arrington Research) was used to monitor subjects' eye positions at 60Hz, after getting calibrated at the beginning of each experimental run. Kruskal-Wallis tests were used to detect systematic differences in the distribution of eye position and movement. The distribution of eye position during search for *communication* and *locomotion* tasks were examined. We find that the distribution of eye position is not affected by search task ($p$=0.17), or by target presence or absence ($p$=0.74), and no significant interactions are present between these two factors ($p$=0.60). To test whether eye movement is affected by target or distractor detection, the distribution of eye position during a 1 sec window around target onset and target offset was studied. The eye position distribution is not affected by target onset ($p$=0.73) or offset ($p$=0.17), and there is no significant interaction between the aforementioned factors ($p$=0.83). Furthermore, the moving-average standard deviation of eye position was studied in a 200ms window to determine systematic differences in rapid moment-to-moment variations in eye position across the two search tasks. There are no significant effects of search task ($p$=0.11), target presence or absence ($p$=0.32), target onset

($p$=0.49), or target offset ($p$=0.36), and there are no significant interactions between these factors ($p$=0.16). Finally, moving-average standard deviation of eye position was included in the model as a nuisance regressor and was regressed out of the BOLD responses.

#### Model estimation and testing

Linearized models were fit in each voxel to estimate model weights that map each set of features (i.e., category, motion-energy, or STIP features) to the measured BOLD responses in each search task in individual subjects. Spatially-informed regularized linear regression (Çelik et al., 2019) with separate regularization terms across feature ($l_f$) and neighbourhood ($l_n$) dimensions was used to fit the models. To capture the hemodynamic response, delayed feature time-courses were concatenated. Delays of two, three, and four samples, corresponding to 4, 6, and 8secs were used. To account for potential correlations between target detection and BOLD responses, a nuisance target-presence regressor was included in the model. The target-presence regressor contained the category regressor for *communication* during search for *communication* task and the category regressor for *locomotion* during search for *locomotion* task. Model fitting for the two search tasks was performed concurrently by concatenating the features and BOLD responses across search tasks (Fig. 2 in the main text). This procedure ensured consistency between the assigned regularization parameters across search tasks and enabled utilisation of the target regressor (Shahdloo et al., 2020).

A nested cross-validation (CV) procedure was used to choose the regularization parameters and estimate model weights. Data from the main experiment were segmented into 60 30-sec blocks. In each of the 10 outer folds, 4 randomly chosen blocks were held-out as validation data. Then, in each of the 10 inner folds, 54 randomly chosen blocks were used as training data and the 2 remaining blocks were used as test data. To fit models for the passive-viewing data, data were segmented into 144 50-sec blocks. In each fold, 8 randomly chosen blocks were held-out as validation data, 132 randomly chosen blocks were used as training data and the 4 remaining blocks were used as test data. We used 10 regularization parameters across the feature dimension in the range $l_f \in [2^5, 2^{17}]$. Similarly, 10 regularization parameters across the neighbourhood dimension in the range $l_f \in [2^{10}, 2^{20}]$ were used. Training data were used to fit models for each ($l_f$, $l_n$) pair independently. Model weights were then used to predict responses in the test data and prediction scores of the fit models were assessed. Prediction scores were taken as product-moment correlation coefficient between measured and predicted voxel responses. The pair of ($l_f$, $l_n$) maximizing the average prediction score across inner CV folds was chosen in each voxel. Finally, the optimal pair of parameters were used to fit models on the union of training and test data in each outer fold and model weights were averaged across the outer folds.

Finally, prediction performance of the fit models were evaluated. In each outer fold, after discarding the nuisance regressors, responses were predicted for the validation data using the fit models and prediction scores were averaged across the search tasks. Prediction scores were then averaged across the outer folds.

#### Variance partitioning

Object-action categories can be correlated with low-level visual features of natural movies (Lescroart and Gallant, 2019), and there is evidence for representation of intermediate-level action features (e.g., action kinematics) across cortex (Jastorff et al., 2010). Therefore, there is a possibility that the estimated category responses are confounded by selectivity for low- and intermediate-level scene features. To control for potential confounds, we performed a variance partitioning analysis. This analysis estimates the response variance that is uniquely explained by the category model after accounting for variance that can be attributed to low- and intermediate-level features captured by the motion-energy and STIP models. To do this, we separately measured the variance explained when all three models (category, motion-energy, and STIP) are fit simultaneously (i.e., combined model), and variance explained when only motion-energy and STIP models are fit simultaneously (i.e., control model). The explained variance ($R^2$) was calculated as squared prediction scores, separately for the combined and control models. Note that from a model fitting perspective, negative prediction scores correspond to zero explained variance. Finally, unique variance explained by the category model was calculated as

$$\widehat{R^2}_{cat} = R^2_{comb} - R^2_{cont} \quad (1)$$

Here $\widehat{R^2}_{cat}$ is the variance uniquely explained by the category model after accounting for low- and intermediate-level features, $R^2_{comb}$ is the variance explained by the combined model, and $R^2_{cont}$ is the variance explained by the union of motion-energy and STIP models in each voxel.

#### Action clustering

To visualise the distribution of actions across the semantic space, the 109 action categories in the movies were separated into distinct clusters. Action content for each short clip was taken as the number of frames where each of the 109 actions were present normalized by the total number of clip frames. This yielded a 109-dimensional action content vector for each clip. The action content vectors were then projected onto the semantic space, and the projections were collapsed into 10 clusters using k-means. The number of clusters was optimized using the elbow method (Thorndike, 1953). To label the clusters, five clips with highest average action contents within each cluster were selected. Four candidate labels for each cluster were then manually assigned and 15 evaluators were asked to score (from 1 to 5) the correspondence of the selected clips to each of the four candidate labels. Finally, the label with the highest score was selected to represent each cluster.

### Supplementary Figures

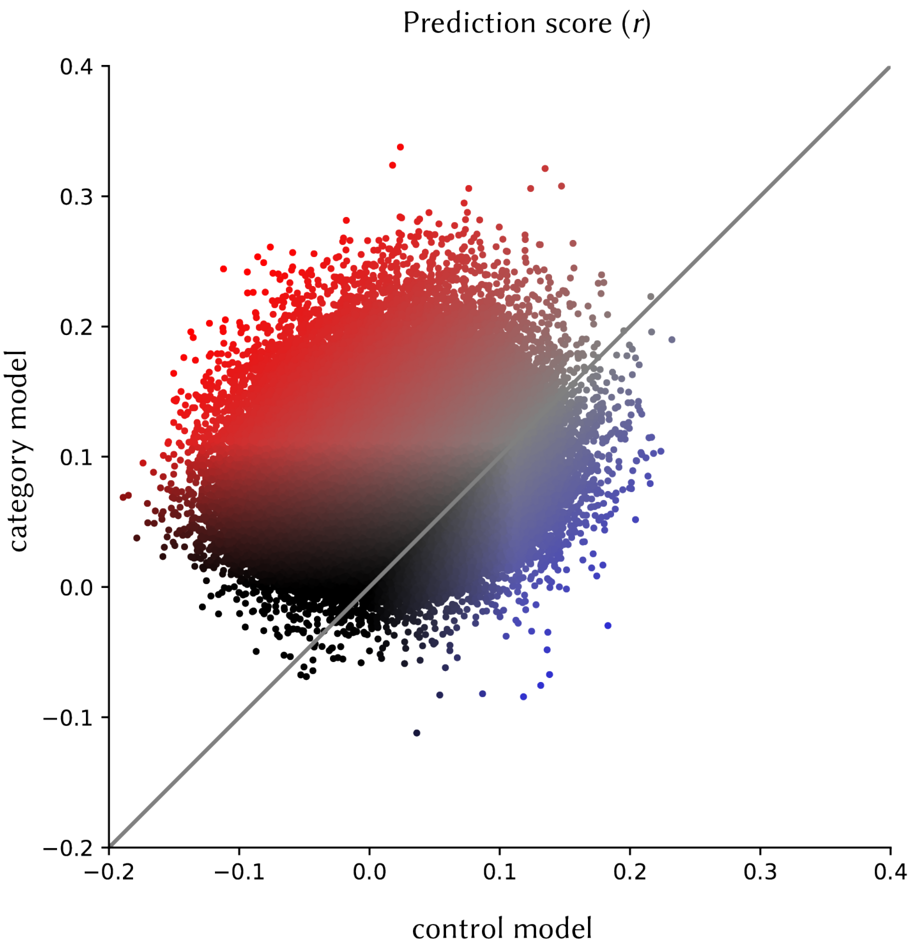


**Supplementary Figure 1. Comparison of category and control models.** The prediction scores (raw product-moment correlation coefficient) of the category and control (the collection of motion-energy and STIP regressors) models were measured for all cortical voxels. Voxels across all subjects are displayed. Each voxel is represented with a dot. Red versus blue dots indicate whether the category model or the control model yields higher prediction scores. Black dots indicate voxels where none of the models has high prediction scores. The category model outperforms the control model in 87.15±2.81% of cortical voxels (mean±sem; average over five subjects).

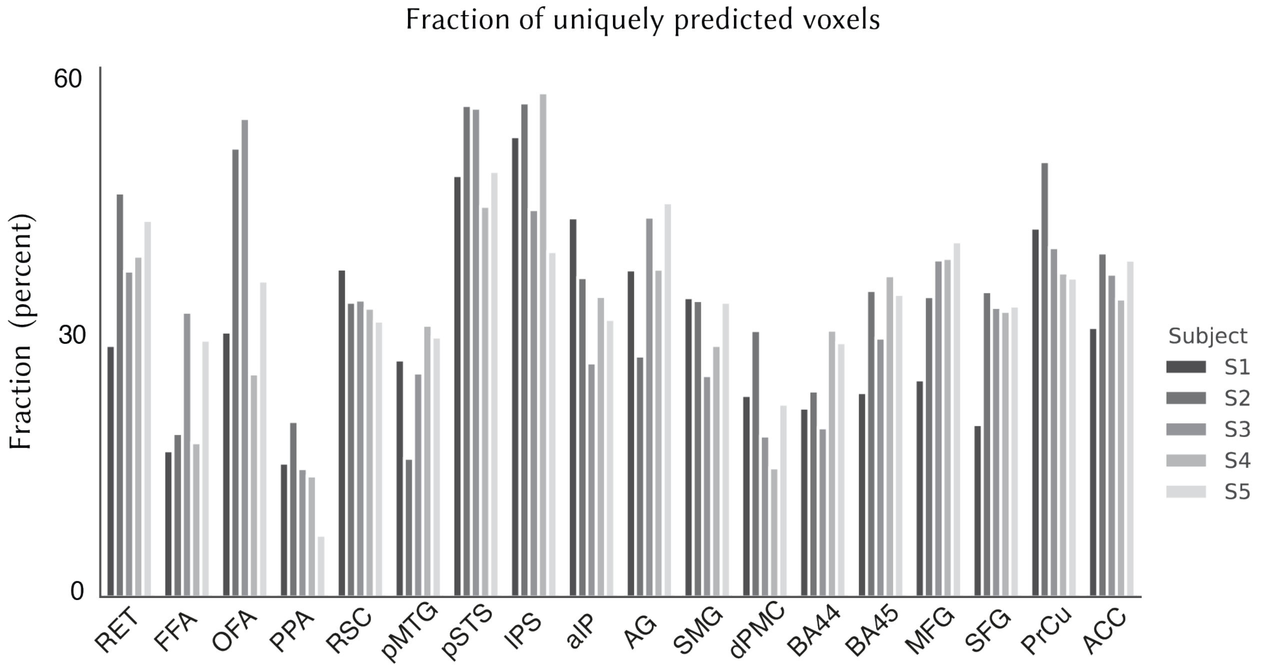


**Supplementary Figure 2. Proportion of uniquely predicted voxels in regions of interest (ROIs).** We identified voxels in which the category model explained unique response variance after accounting for low-level motion-energy, and intermediate-level STIP stimulus features by performing a variance partitioning analysis (see *Supplementary Methods*). Fraction of these *semantic voxels* is shown across ROIs, in individual subjects.

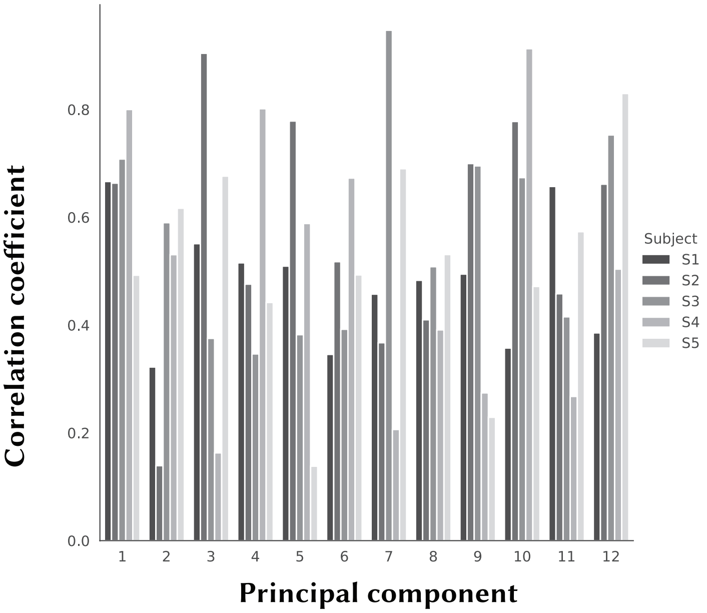


**Supplementary Figure 3. Consistency of the semantic space across subjects.** To test whether the estimated semantic space is consistent across subjects, leave-one-out cross-validation was performed. In each cross-validation fold, best-predicted voxels from four subjects were used to derive 12 PCs to construct a semantic space. In the left-out subject, semantic tuning profile for each voxel was obtained by projecting action category responses during passive viewing onto the derived PCs. Next, product-moment correlation coefficient was calculated between the tuning profiles in the derived space and the tuning profiles in the original semantic space. Results were averaged across semantic voxels in the left-out subject. Correlation coefficients are shown for each PC and each subject. The cross-validated semantic spaces consistently correlate with the original semantic space.

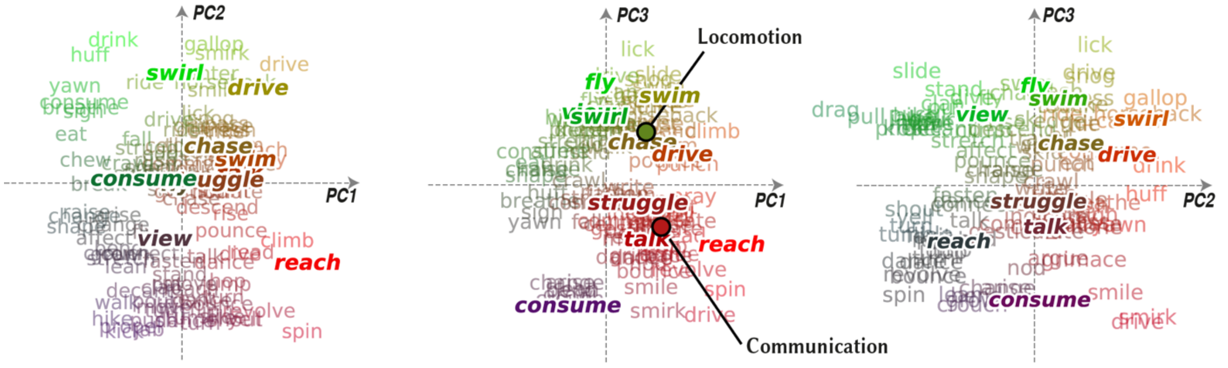


**Supplementary Figure 4. Distribution of action categories across PCs.** To illustrate the distribution of action categories embedded within the semantic space, action categories were projected onto the PCs. Projections onto the first three PCs are shown (words in regular font show projections of individual categories). To facilitate illustration, categories were collapsed into 10 clusters and cluster centres were also projected onto the PCs (bold-italic words; see *Supplementary Methods*). Average location of the *communication* and *locomotion* actions are specified with red and green dots. The estimated semantic space captures reasonable semantic variance across action categories in natural movies.

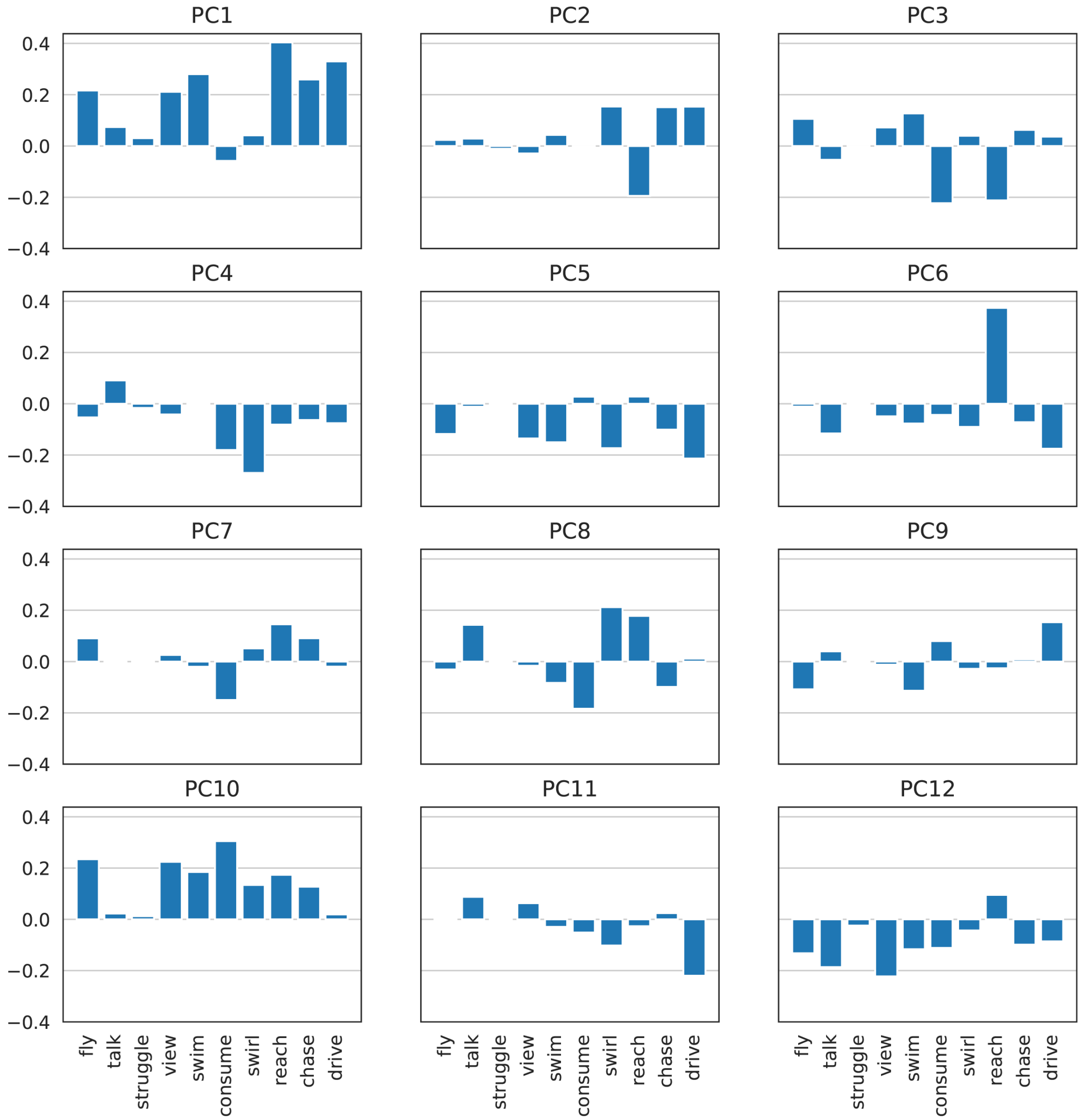


**Supplementary Figure 5. Projections of action category clusters onto PCs.** Each of the 109 action categories were projected onto the twelve semantic principal components (PCs). The projections were then clustered into 10 groups using k-means and labelled for interpretation (see *Supplementary Methods*). The projections of the cluster centres onto 12 PCs are shown. The first three dimensions were used to visualise the semantic space (Supp. Fig. 3). The first dimension distinguishes between self-movements (e.g., swirl, consume) and actions that are targeted toward other humans or objects (e.g., reach, talk). The second dimension distinguishes between dynamic (e.g., drive, chase) versus static actions (e.g., consume, struggle). The third dimension distinguishes between actions that involve humans (e.g., talk, reach) and dynamic actions (e.g., fly, swirl).

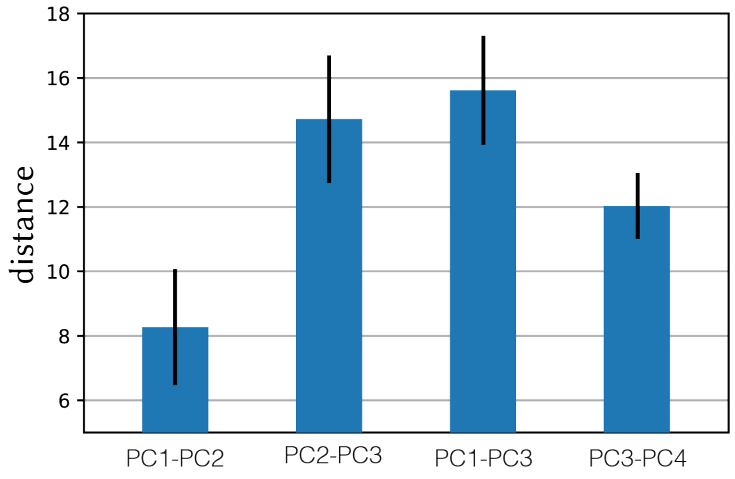


**Supplementary Figure 6. The distance between target actions in subspaces spanned by different pairs of PCs.** To visualise attentional modulation of semantic representation in Fig. 4 of the main text, we compared projections of action category responses onto a pair of PCs across the search tasks. To maximize our sensitivity in visualising the attentional modulations, we chose the pair of dimensions that maximally separates the actions belonging to the two target categories (i.e., *communication* and *locomotion* categories). The Mahalanobis distance between communication actions and locomotion actions (mean±sem across communication and locomotion actions) in the subspace spanned by each pair of PCs is shown. Target actions are maximally separated across the subspace spanned by the first and third PCs.

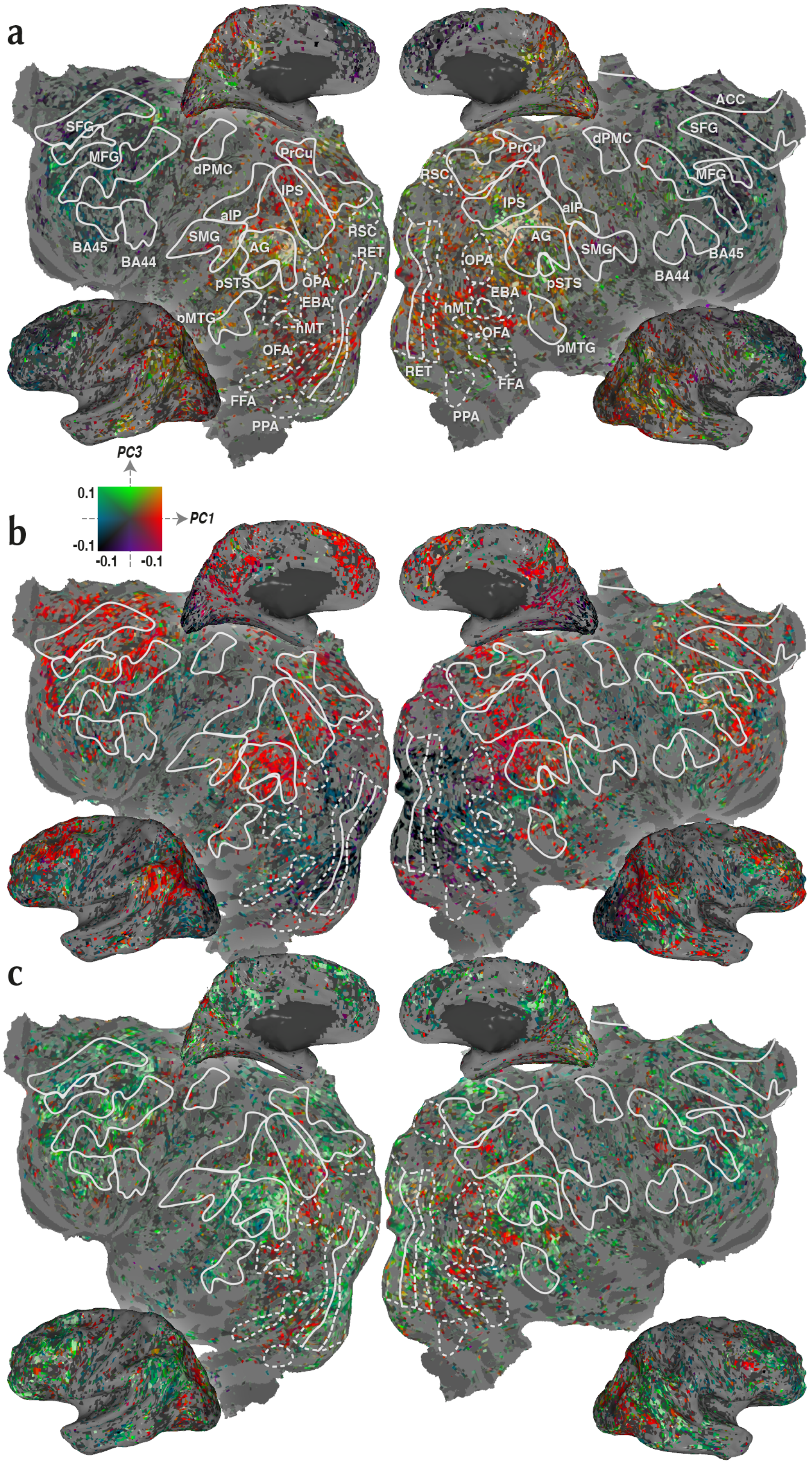


**Supplementary Figure 7. Cortical flat maps of semantic representation for subject S1.** Action category responses during **a.** passive viewing, **b.** search for *communication*, and **c.** search for *locomotion* categories were projected onto the semantic space in subject S1. A two-dimensional colourmap was used to colour each voxel based on the projection values along the first and third semantic dimensions (see colour legend). Voxels where the category model does not explain unique response variance after accounting for low- and intermediate-level stimulus features are masked (bootstrap test, $q$(FDR)<0.05). Regions of interest are illustrated by white borders. Several important sulci are illustrated by dashed grey lines. Abbreviations for regions of interest and sulci are listed in *Supplementary Methods*. Many voxels across occipitotemporal, parietal, and prefrontal cortices shift their tuning toward targets.

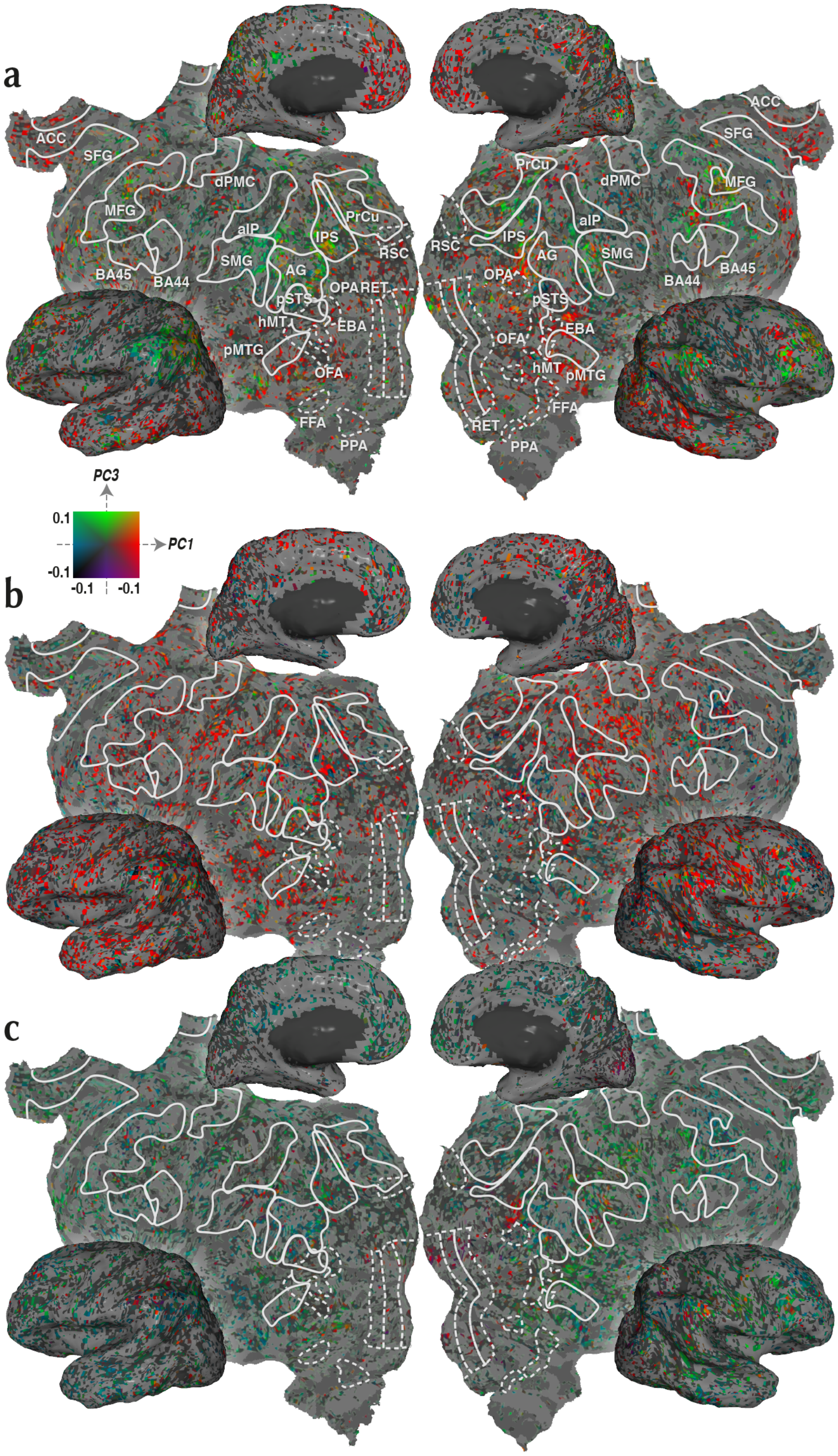


**Supplementary Figure 8. Cortical flat maps of semantic representation for subject S2.** Action category responses during **a.** passive viewing, **b.** search for *communication*, and **c.** search for *locomotion* categories were projected onto the semantic space in subject S2. Formatting is identical to Supp. Fig. 7.

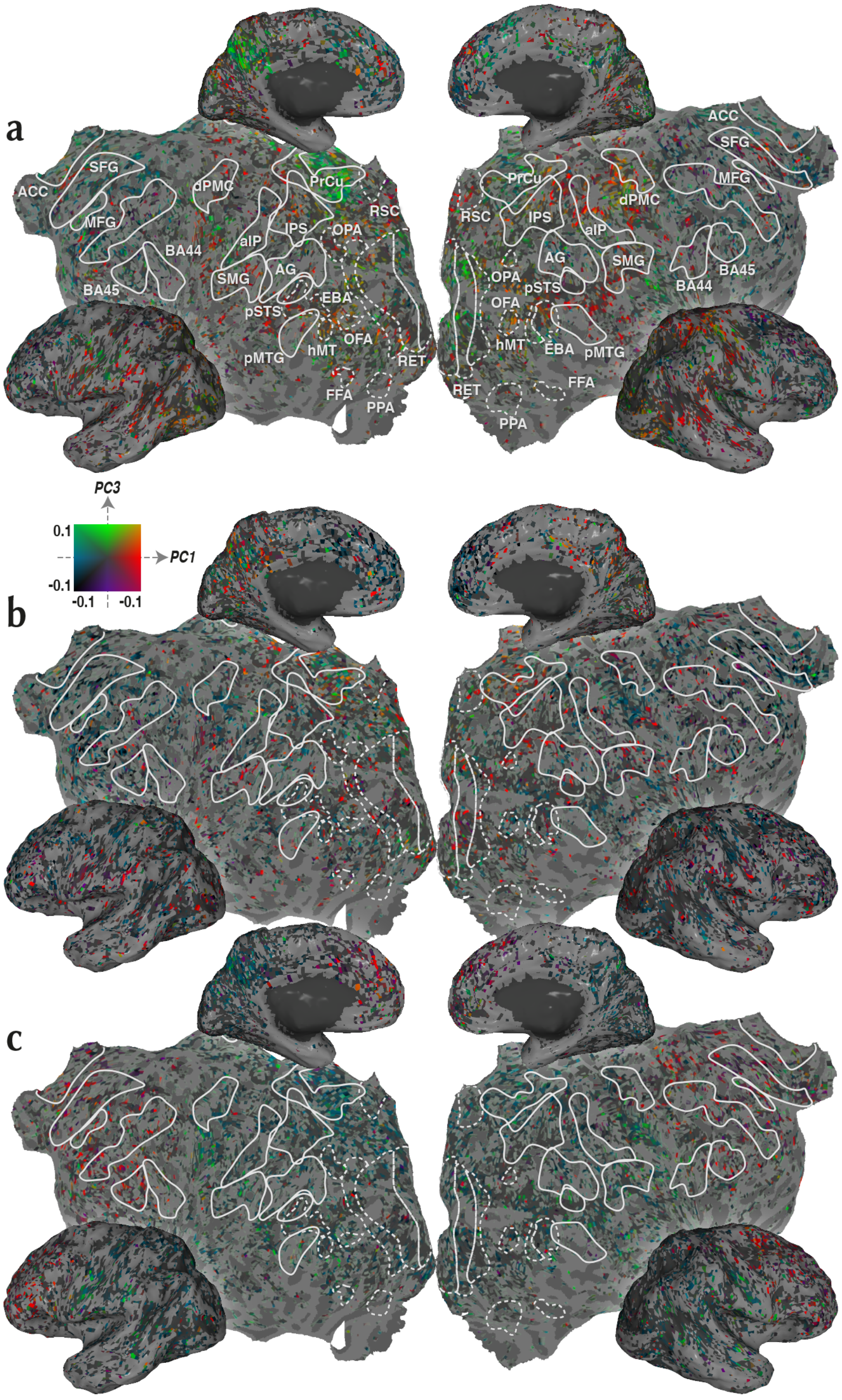


**Supplementary Figure 9. Cortical flat maps of semantic representation for subject S3.** Action category responses during **a.** passive viewing, **b.** search for *communication*, and **c.** search for *locomotion* categories were projected onto the semantic space in subject S3. Formatting is identical to Supp. Fig. 7.

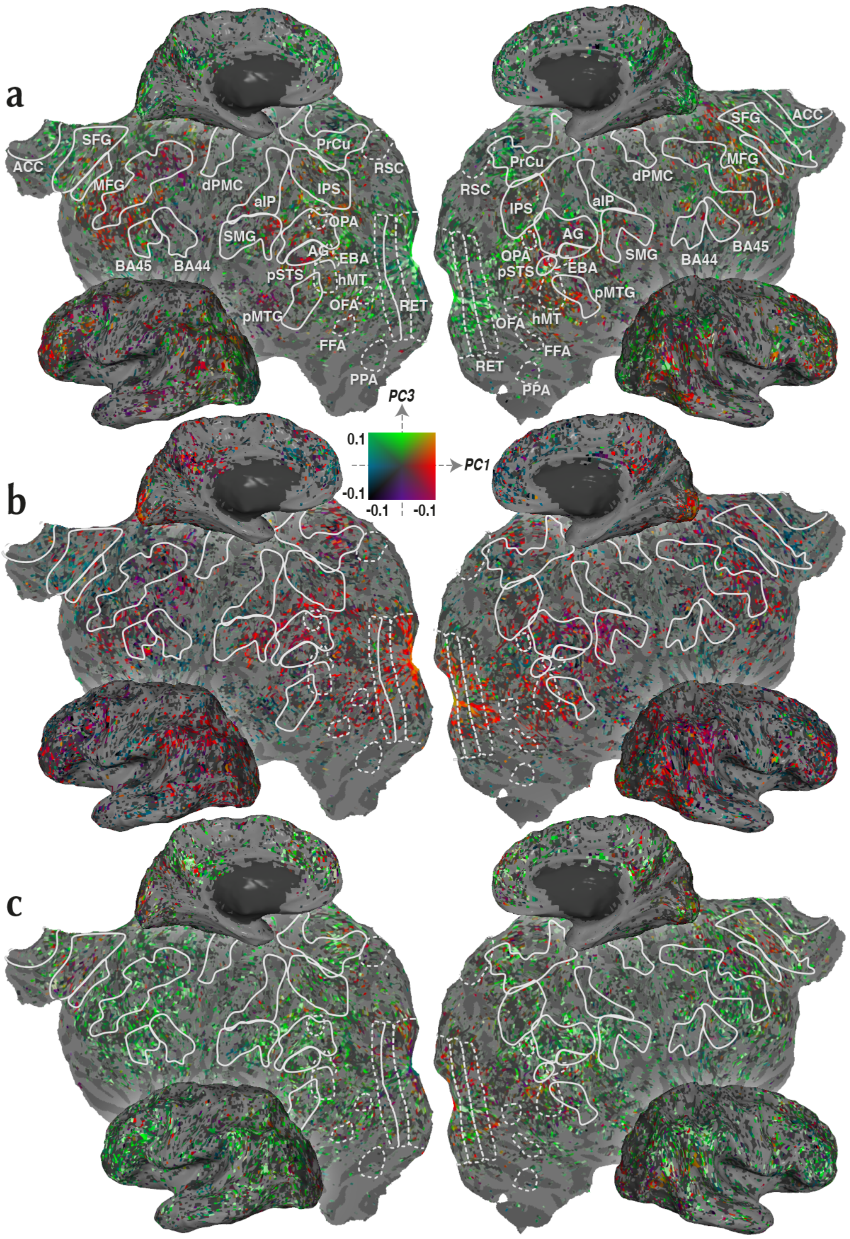


**Supplementary Figure 10. Cortical flat maps of semantic representation for subject S4.** Action category responses during **a.** passive viewing, **b.** search for *communication*, and **c.** search for *locomotion* categories were projected onto the semantic space in subject S4. Formatting is identical to Supp. Fig. 7.

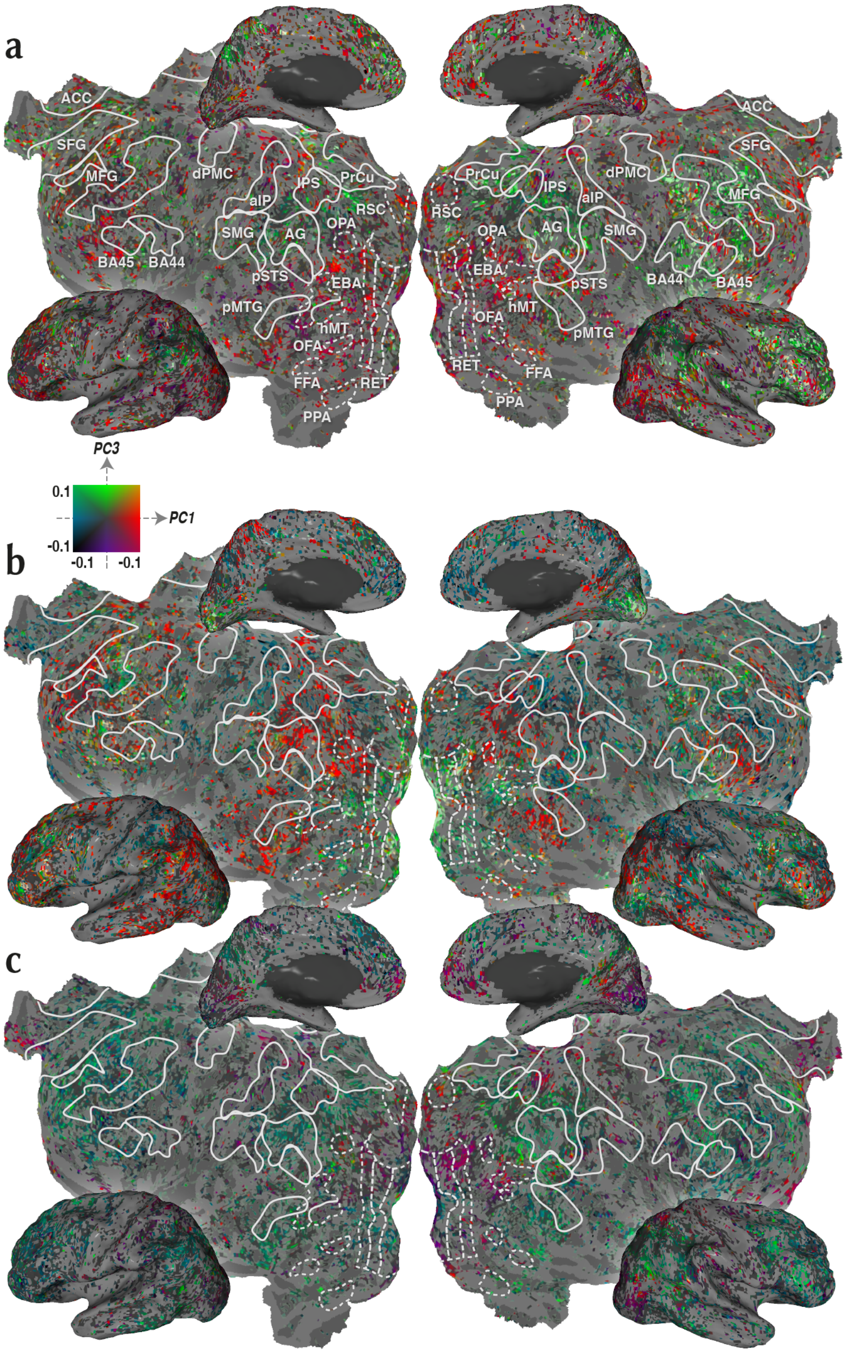


**Supplementary Figure 11. Cortical flat maps of semantic representation for subject S5.** Action category responses during **a.** passive viewing, **b.** search for *communication*, and **c.** search for *locomotion* categories were projected onto the semantic space in subject S5. Formatting is identical to Supp. Fig. 7.

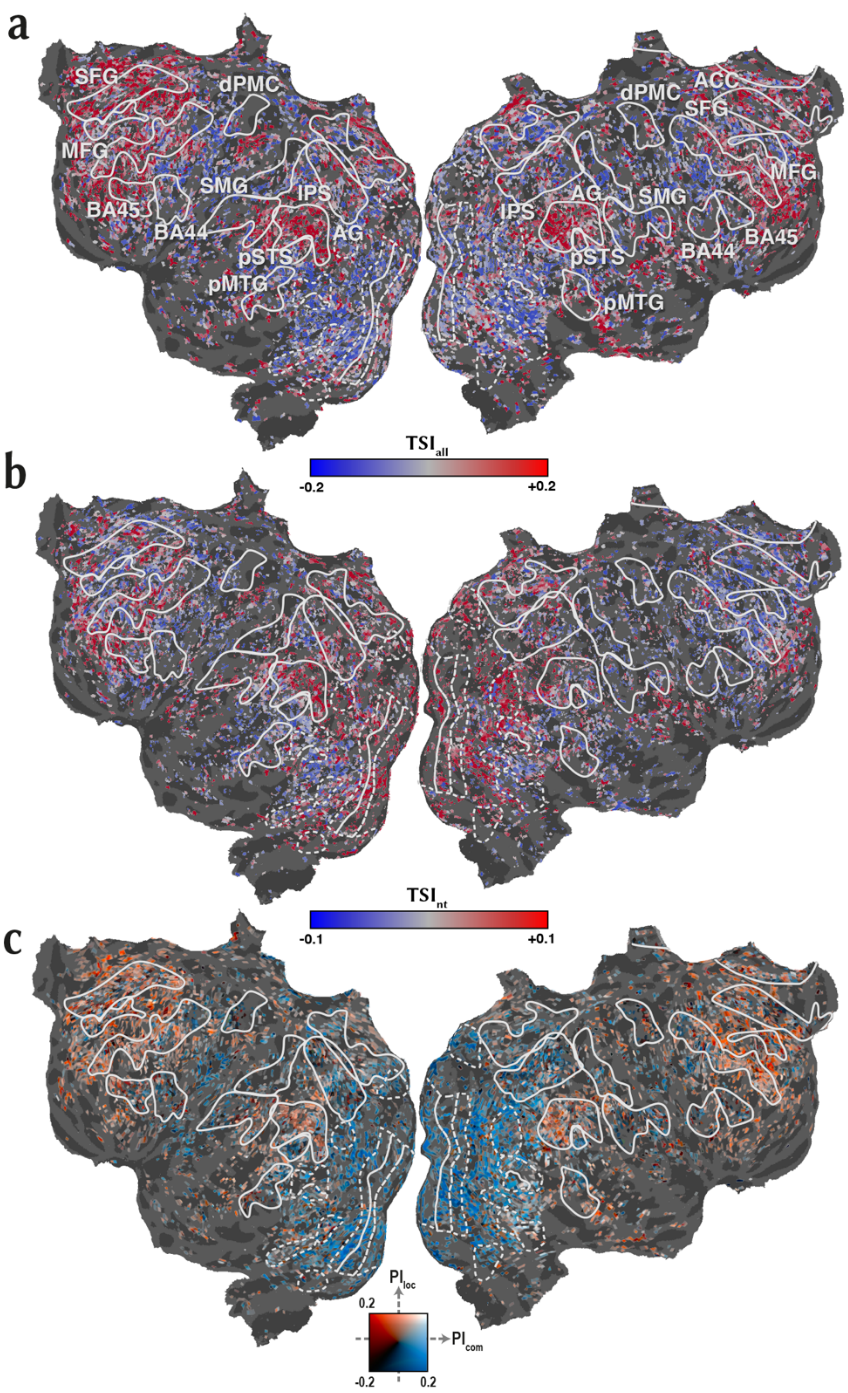


**Supplementary Figure 12. Cortical flat maps of TSI and PI for subject S1. a.** Tuning shift index for all action categories ($TSI_{all}$), **b.** tuning shift for nontarget categories ($TSI_{nt}$), and **c.** preference index values ($PI_{com}$, $PI_{loc}$) were projected onto the semantic space in subject S1 (see legends in Fig. 5 and 6 in the main text for the colour map). Only significant voxels are shown (bootstrap test, $p<0.05$). Formatting is identical to Supp. Fig. 7.

a

b

c

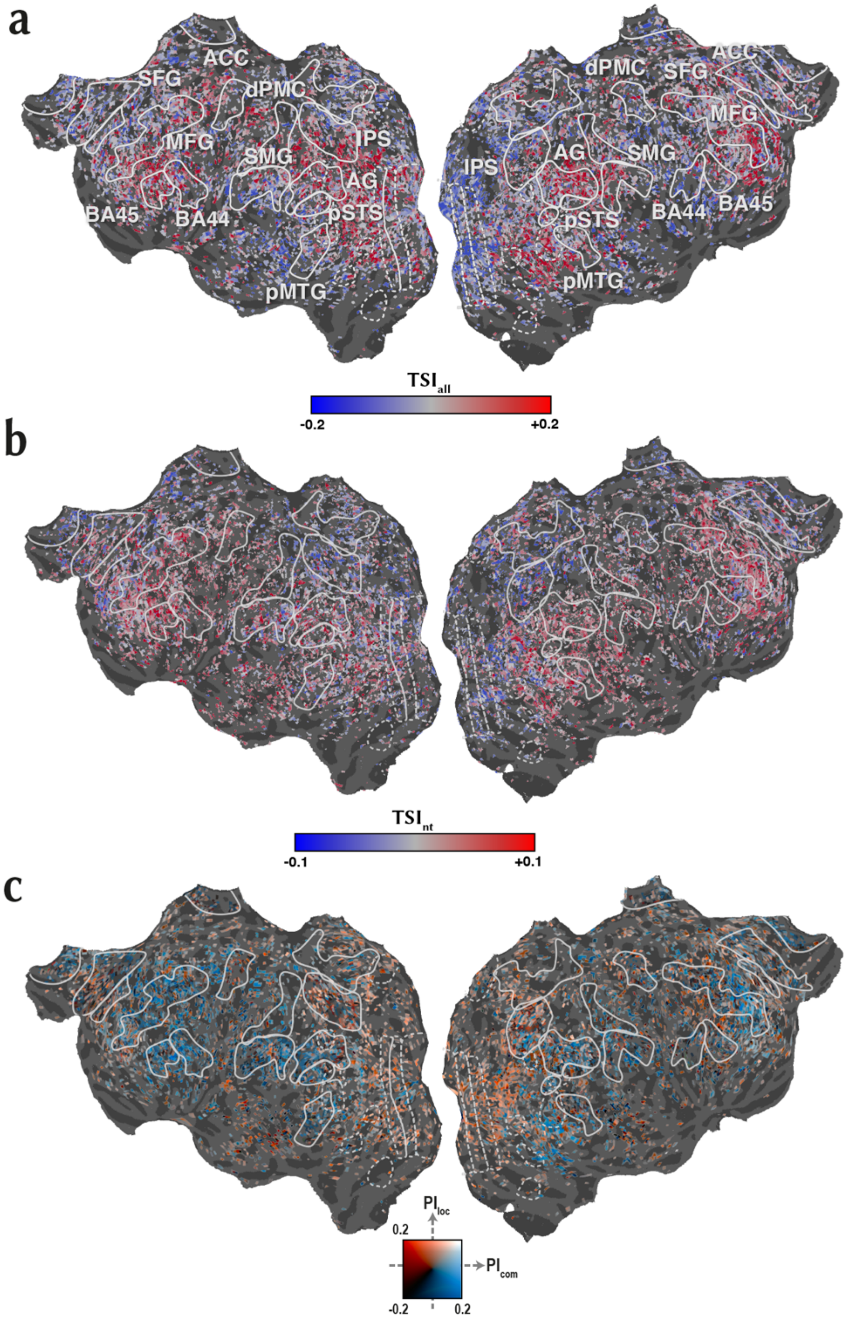


**Supplementary Figure 13. Cortical flat maps of TSI and PI for subject S2. a.** Tuning shift index for all action categories ($TSI_{all}$), **b.** tuning shift for nontarget categories ($TSI_{nt}$), and **c.** preference index values ($PI_{com}$, $PI_{loc}$) were projected onto the semantic space in subject S2 (see legends in Fig. 5 and 6 in the main text for the colour map). Only significant voxels are shown (bootstrap test, $p<0.05$). Formatting is identical to Supp. Fig. 7.

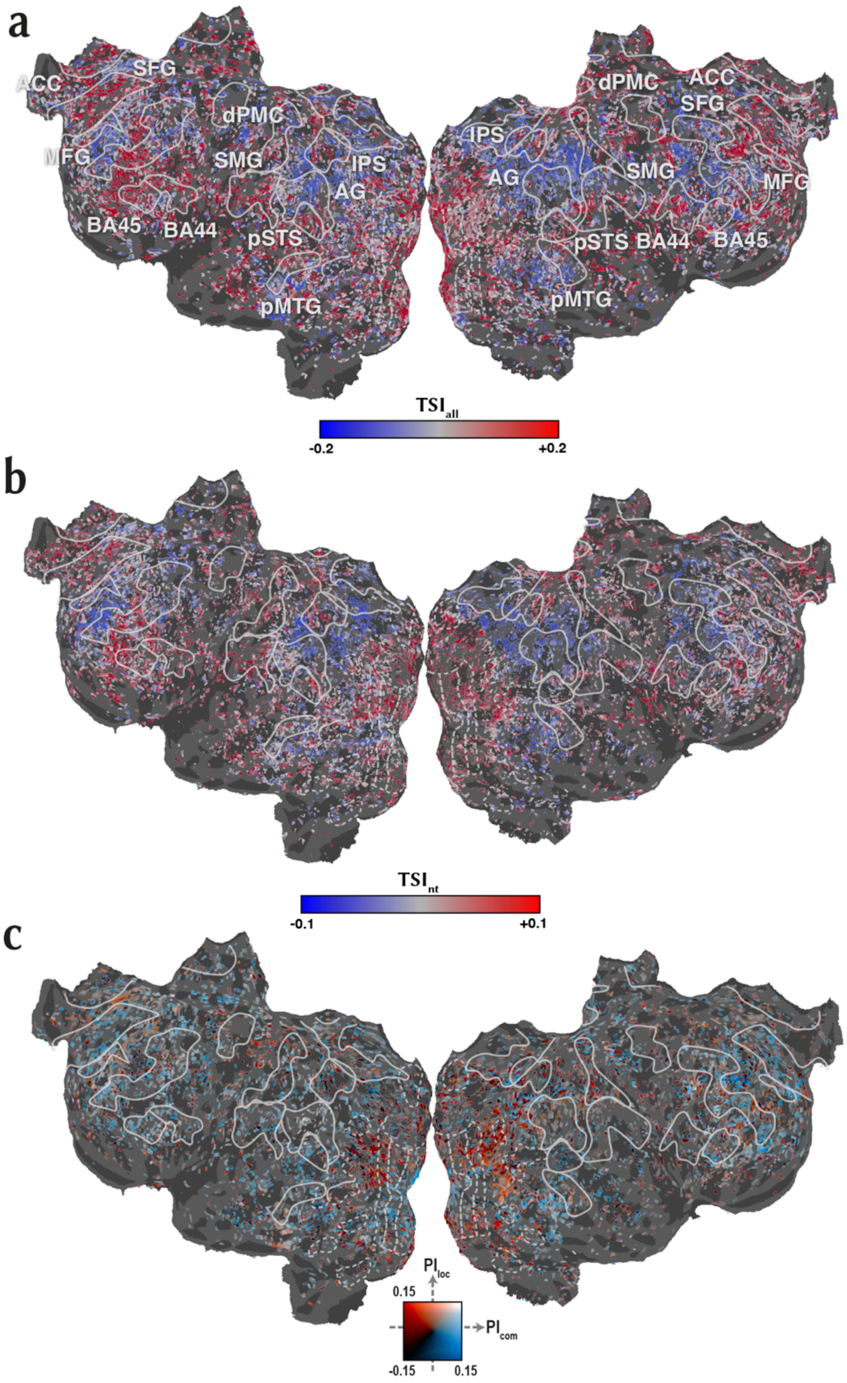


**Supplementary Figure 14. Cortical flat maps of TSI and PI for subject S3. a.** Tuning shift index for all action categories ($TSI_{all}$), **b.** tuning shift for nontarget categories ($TSI_{nt}$), and **c.** preference index values ($PI_{com}$, $PI_{loc}$) were projected onto the semantic space in subject S3 (see legends in Fig. 5 and 6 in the main text for the colour map). Only significant voxels are shown (bootstrap test, $p<0.05$). Formatting is identical to Supp. Fig. 7.

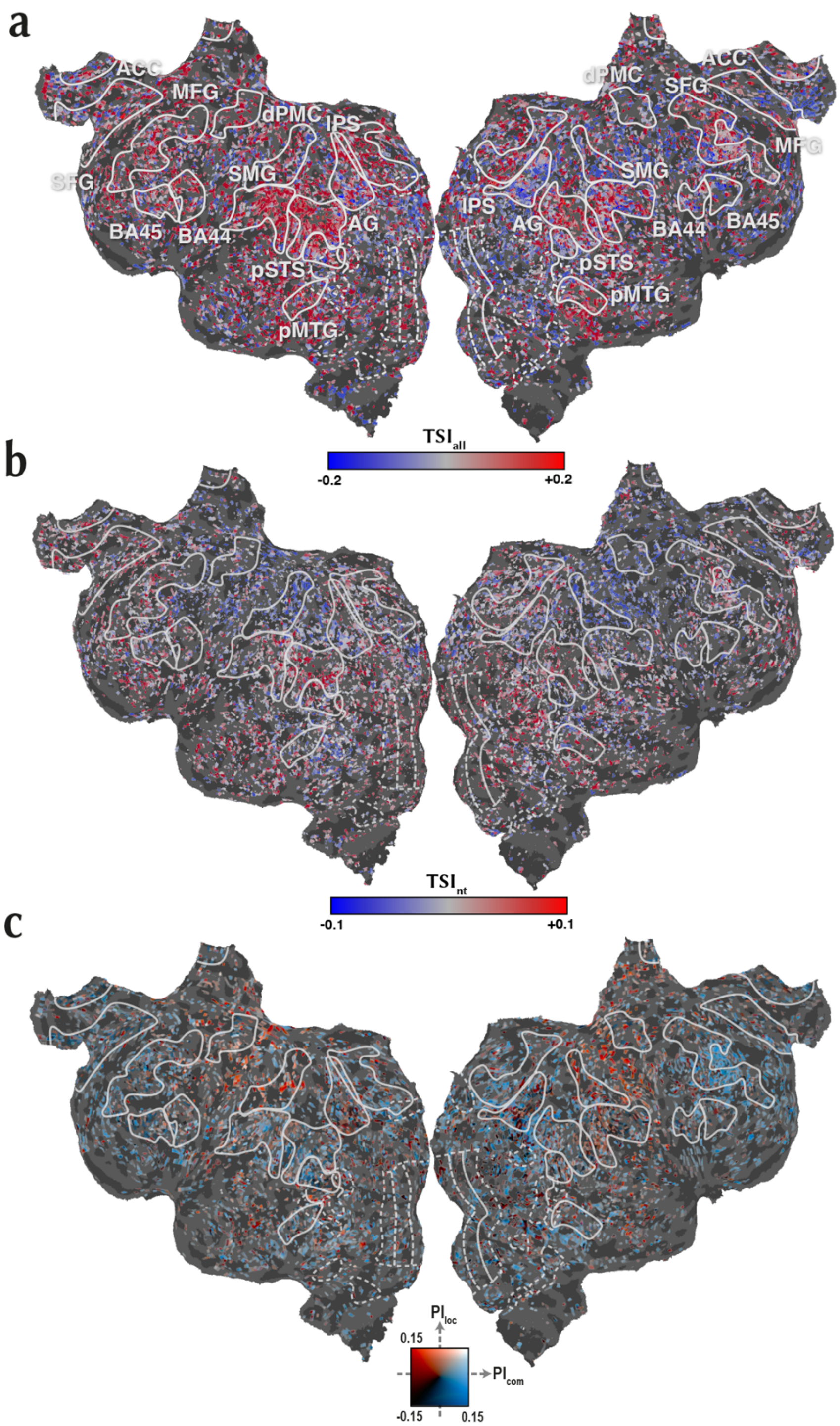


**Supplementary Figure 15. Cortical flat maps of TSI and PI for subject S4. a.** Tuning shift index for all action categories ($TSI_{all}$), **b.** tuning shift for nontarget categories ($TSI_{nt}$), and **c.** preference index values ($PI_{com}$, $PI_{loc}$) were projected onto the semantic space in subject S4 (see legends in Fig. 5 and 6 in the main text for the colour map). Only significant voxels are shown (bootstrap test, $p$<0.05). Formatting is identical to Supp. Fig. 7.

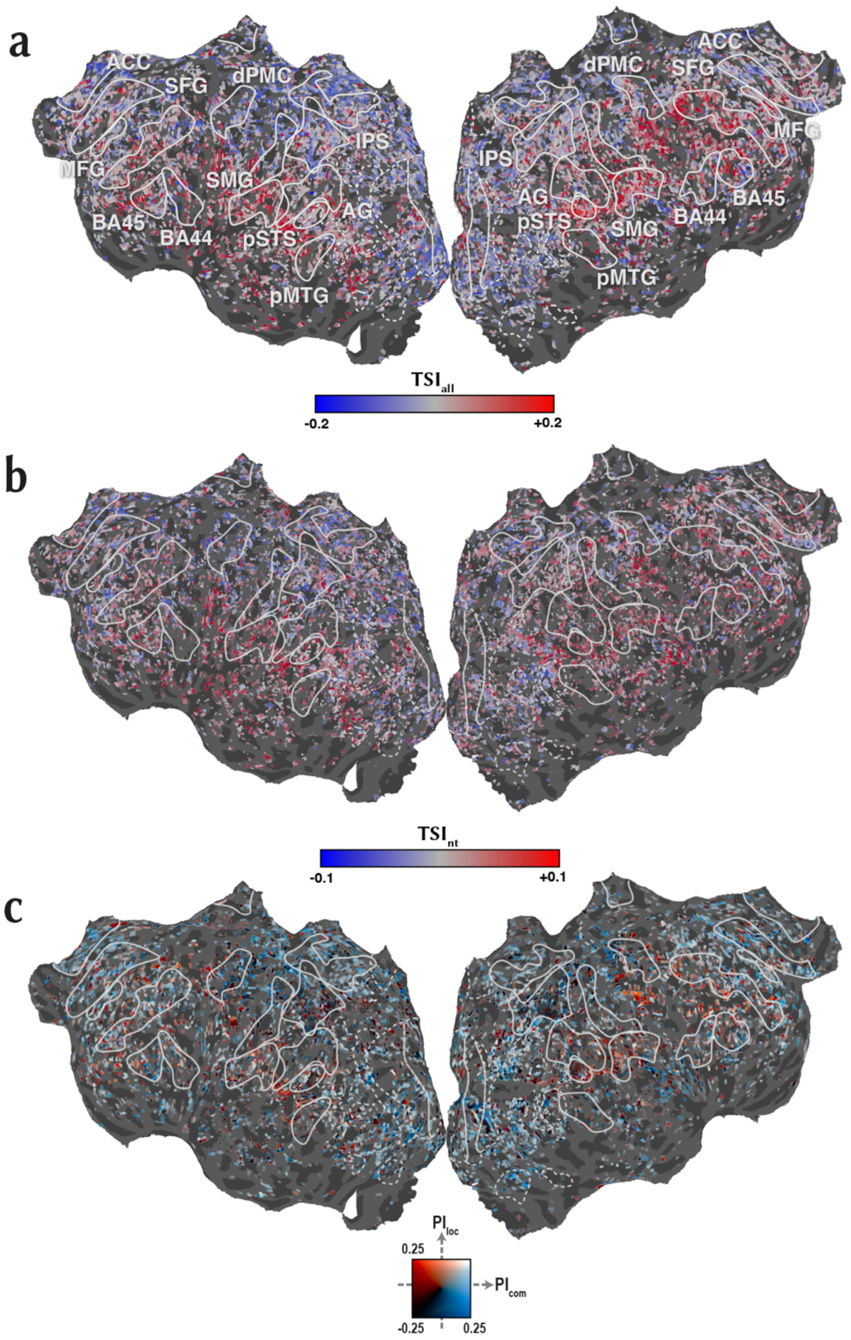


**Supplementary Figure 16. Cortical flat maps of TSI and PI for subject S5. a.** Tuning shift index for all action categories ($TSI_{all}$), **b.** tuning shift for nontarget categories ($TSI_{nt}$), and **c.** preference index values ($PI_{com}$, $PI_{loc}$) were projected onto the semantic space in subject S5 (see legends in Fig. 5 and 6 in the main text for the colour map). Only significant voxels are shown (bootstrap test, $p<0.05$). Formatting is identical to Supp. Fig. 7.

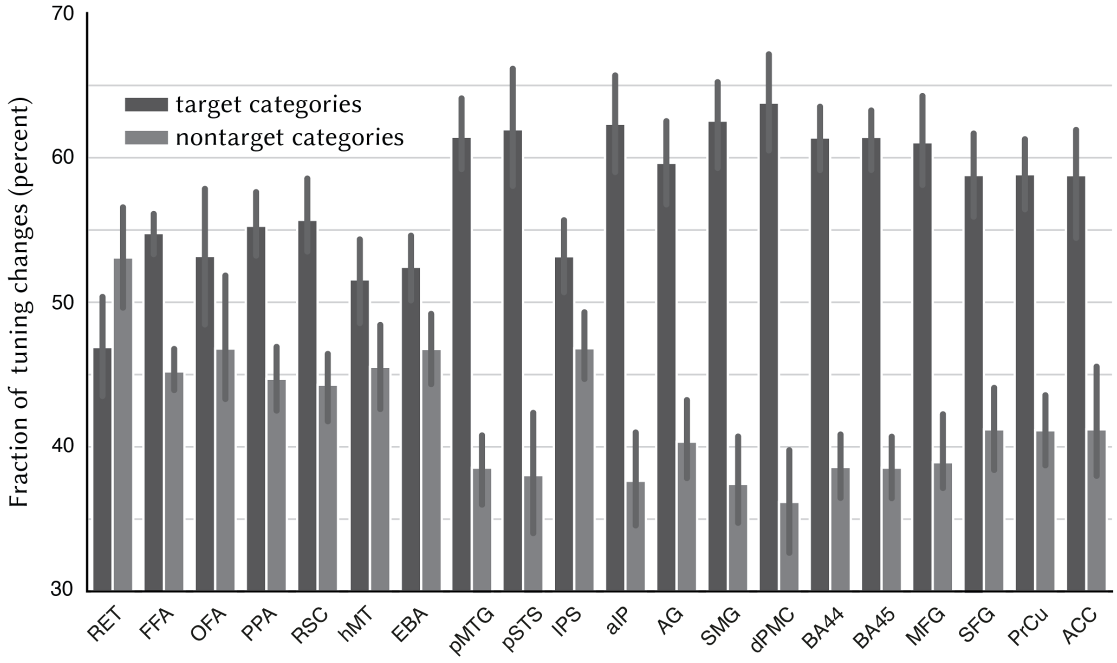


**Supplementary Figure 17. Fraction of the overall tuning shifts.** Fraction of the overall tuning shifts explained by shifts in tuning for target categories (mean±sem across subjects) and nontarget categories (i.e., excluding the union of communication and locomotion categories) is shown. Target categories explain a greater portion of the overall tuning shifts broadly across ROIs, except for early retinotopic areas. At the same time, nontarget categories significantly contribute to the overall tuning shifts.

### Supplementary References


Emin Çelik, Salman Ul Hassan Dar, Özgür Yılmaz, Ümit Keleş, and Tolga Çukur. Spatially informed voxelwise modeling for naturalistic fMRI experiments. *NeuroImage*, 186: 741–757, February 2019.

Anders M Dale, Bruce Fischl, and Martin I Sereno. Cortical surface-based analysis: I. Segmentation and surface reconstruction. *NeuroImage*, 9 (2): 179–194, January 1999.

Christophe Destrieux, Bruce Fischl, Anders Dale, and Eric Halgren. Automatic parcellation of human cortical gyri and sulci using standard anatomical nomenclature. *NeuroImage*, 53 (1): 1–15, October 2010.

Karl J Friston, C D Frith, R Turner, and R S J Frackowiak. Characterizing evoked hemodynamics with fMRI. *NeuroImage*, 2 (2): 157–165, January 1995.

James S Gao, Alexander G Huth, Mark D Lescroart, and Jack L Gallant. PyCortex: an interactive surface visualiser for fMRI. *Frontiers in Neuroinformatics*, 9 (22): 162, September 2015.

Alexander G Huth, Shinji Nishimoto, An T Vu, and Jack L Gallant. A Continuous Semantic Space Describes the Representation of Thousands of Object and Action Categories across the Human Brain. *Neuron*, 76 (6): 1210–1224, December 2012.

Jan Jastorff, Chiara Begliomini, Maddalena Fabbri-Destro, Giacomo Rizzolatti, and Guy A Orban. Coding observed motor acts: Different organisational principles in the parietal and premotor cortex of humans. *Journal of Neurophysiology*, 104 (1): 128–140, July 2010.

Mark D Lescroart and Jack L Gallant. Human Scene-Selective Areas Represent 3D Configurations of Surfaces. *Neuron*, 101 (1): 178–192.e7, January 2019.

Martin Reuter, Nicholas J Schmansky, H Diana Rosas, and Bruce Fischl. Within-subject template estimation for unbiased longitudinal image analysis. *NeuroImage*, 61 (4): 1402–1418, July 2012.

Mohammad Shahdloo, Emin Çelik, and Tolga Çukur. Biased Competition in Semantic Representation During Natural Visual Search. *NeuroImage*, page 116383, November 2019.

Stephen M Smith. Fast robust automated brain extraction. *Human Brain Mapping*, 17 (3): 143–155, November 2002.

Thorndike, R.L. Who belongs in the family?. *Psychometrika* 18, 267–276, 1953.

Timothy D Verstynen and Vibhas Deshpande. Using pulse oximetry to account for high and low frequency physiological artifacts in the BOLD signal. *NeuroImage*, 55 (4): 1633–1644, April 2011.
